# Actomyosin Dynamics Determine the Extension and Retraction of Filopodia on Neuronal Dendrites

**DOI:** 10.1101/057919

**Authors:** Olena O. Marchenko, Sulagna Das, Ji Yu, Igor L. Novak, Vladimir I. Rodionov, Nadia Efimova, Tatiana Svitkina, Charles W. Wolgemuth, Leslie M. Loew

**Author notes:** Olena O. Marchenko R.D.Berlin Center for Cell Analysis and Modeling Uconn Health Center 400 Farmington Ave, Farmington, CT 06030. Sulagna Das Department of Anatomy and Structural Biology, Albert Einstein College of Medicine 1300 Morris Park Ave, Bronx, NY 10461, New York, NY. Ji Yu R.D.Berlin Center for Cell Analysis and Modeling Uconn Health Center 400 Farmington Ave, Farmington, CT 06030. Igor L. Novak R.D.Berlin Center for Cell Analysis and Modeling Uconn Health Center 400 Farmington Ave, Farmington, CT 06030. Vladimir I. Rodionov R.D.Berlin Center for Cell Analysis and Modeling Uconn Health Center 400 Farmington Ave, Farmington, CT 06030. Nadia Efimova Department of Biology University of Pennsylvania Philadelphia, PA. Tatiana Svitkina Department of Biology University of Pennsylvania Philadelphia, PA. Charles W. Wolgemuth Departments of Physics and Molecular and Cellular Biology University of Arizona Tucson, AZ 85721. Leslie M. Loew R.D. Berlin Center for Cell Analysis and Modeling Uconn Health Center Farmington, CT 06030.

## Abstract

**Impact Statement:** In this study, using a combination of computational and experimental approaches we show that a complex dynamic behavior of dendritic filopodia that is essential for synaptogenesis is explained by an interplay among forces generated by actin retrograde flow, myosin contractility, and substrate adhesion.

**Abstract:** Dendritic filopodia are actin-filled dynamic subcellular structures that sprout on neuronal dendrites during neurogenesis. The exploratory motion of the filopodia is crucial for synaptogenesis but the underlying mechanisms are poorly understood. To study the filopodial motility, we collected and analyzed image data on filopodia in cultured rat hippocampal neurons. We hypothesized that mechanical feedback among the actin retrograde flow, myosin activity and substrate adhesion gives rise to various filopodial behaviors. We have formulated a minimal one-dimensional partial differential equation model that reproduced the range of observed motility. To validate our model, we systematically manipulated experimental correlates of parameters in the model: substrate adhesion strength, actin polymerization rate, myosin contractility and the integrity of the putative microtubule-based barrier at the filopodium base. The model predicts the response of the system to each of these experimental perturbations, supporting the hypothesis that our actomyosin-driven mechanism controls dendritic filopodia dynamics.

## Introduction

Formation of synapses during development and their maintenance in the adult organism are required for the proper functioning of neuronal circuitry. Spines, micrometer-scale protrusions from neuronal dendrites, receive the majority of synaptic inputs in the central nervous system. Regulation of spine density, morphology and spatial distribution is required for synaptic stability and plasticity. Abnormal density and morphology are associated with impaired motor and cognitive functions underlying neurological disorders such as autism, schizophrenia and fragile-X syndrome (Segal, 1995, Wilson et al., 2010, Irwin et al., 2000, Lin and Koleske, 2010).

Spine morphogenesis is a delicate process that starts from the formation of dendritic filopodium (Ziv and Smith, 1996, Hotulainen and Hoogenraad, 2010, Korobova and Svitkina, 2010). Dendritic filopodia are dynamic protrusions up to 15μm in length that explore their extracellular environment and connect with the presynaptic axons (Kayser et al., 2008). The population of motile filopodia is highest during early neuron development and gradually decreases as the neuron ages. Starting from the 4^th^ day of *in vitro* (DIV4) hippocampal cell culture, when neurites are fully differentiated into axons and dendrites, filopodia are numerous and highly motile (Ziv and Smith, 1996). They extend 2-10 microns away from the dendritic shaft to establish contact with nearby axons and also can retract back toward the dendrite. The protrusion/retraction cycle repeats until filopodia stabilize or dissolve back into their parent dendrite. By DIV13 most of the filopodia observed in culture are non-motile, whether solitary or connected to an axon. In the same time frame, the first spines start to appear on the dendrites. The initial axon-dendritic contact and its stabilization are considered to be the key events in spinogenesis (Kayser et al., 2008, Hotulainen and Hoogenraad, 2010). However, the dynamics of spine morphogenesis from filopodia has not been fully characterized and the mechanism of dendritic filopodial motility is poorly understood.

What is known is that dendritic filopodial motility is actin-based. The main components of actin network dynamics in motile cells have been extensively studied, resulting in models that describe various mechanisms of leading edge protrusion (Leibler and Huse, 1993, Lammermann and Sixt, 2009, Mogilner and Rubinstein, 2005). From models of keratocyte and nerve growth cone motility we know that actin-based motility arises from the force that polymerizing actin filaments exert on the membrane, which is opposed by membrane tension and substrate adhesion (Medeiros et al., 2006). Myosin II contracts the actin filaments generating the actin retrograde flow that pulls filaments away from the leading edge (Verkhovsky et al., 1995). If the actin filament polymerization rate overcomes the retrograde flow, a local protrusion forms. How are actin polymerization, contraction and local adhesion mechanisms marshaled to control dendritic filopodial motility and stability? Previous models of filopodial dynamics treat the system as a stiff elastic rod compressed by the membrane resistance force without considering adhesion or contractility(Mogilner and Rubinstein, 2005); others have focused on the adhesion to and compliance of the substrate (Chan and Odde, 2008). These earlier studies do not consider the unique molecular composition of dendritic filopodia.

The molecular composition of the actin network in dendritic filopodia, which lack filament bundling proteins found in conventional filopodia (Korobova and Svitkina, 2010), affects the rheological properties making it more viscous compared to bundled actin (Kim et al., 2009). For example, the actin network in dendritic spines is 4-5 times more viscous than the average cell cytoskeleton (Smith et al., 2007). The actin cytoskeleton inside a dendritic filopodium consists of short branched filaments of mixed polarity (Portera-Cailliau et al., 2003, Korobova and Svitkina, 2010), which can make them sensitive to myosin contractile stresses. Adhesion to the substrate can affect actin retrograde flow and govern the retraction phase of the filopodia motility cycle (Yamashiro and Watanabe, 2014). Furthermore, substrate adhesion is required for neurogenesis(Lin and Koleske, 2010) and may be important for dendritic filopodia behavior.

Therefore, we expect the dynamics of dendritic filopodia to differ from that of conventional filopodia. Such a range of behaviors could be involved in the establishment and stabilization of initial synaptic connections in the developing nervous system (Portera-Cailliau et al., 2003). Indeed, in this paper we characterize a wide range of motility states, from non-motile to persistently fluctuating filopodia. To understand these diverse motility behaviors, we develop a minimal mathematical model of dendritic filopodia dynamics that explains the spatiotemporal regulation of actin and myosin dynamics and describes conditions under which dynamic filopodia are maintained. The model employs partial differential equations that balance the forces of substrate adhesion, actin retrograde flow, and myosin contractility, while accounting for actin polymerization and diffusion. To validate the proposed mechanism experimentally, we perturbed key model parameters using various manipulations, comparing the model predictions with the results of motility analysis of dendritic filopodia on hippocampal neurons in culture.

## Results

To collect sufficient data on filopodia dynamics, we performed 2hr (Δt=1s) time-lapse recordings of filopodia extending and retracting from the dendrites of young neurons (DIV4-12) (*Fig.1A*). Filopodial dynamics were then quantified by tracking filopodia length automatically using our custom-written software FiloTracker (*Movie S1A, Fig.1B, C*). Having collected trajectories from 938 filopodia (83 neurons), we classified dendritic filopodia by their motility patterns into filopodia with transient and continual dynamics, and non-motile filopodia (*Fig S1.A–C*). Filopodia with continual dynamics displayed lifetimes t>20min with regular protrusion/retraction cycles(*Fig. S1A*). Filopodia with transient motility were defined by irregular short-lived bursts of motility (*Fig. S1B*), and filopodia with constant length did not exhibit any motility, displaying invariable length for the duration of the recording session (20min or more) (*Fig. S1C*). Filopodia were more motile during early neurodevelopment with a motile fraction of 41%±22 for DIV 3-9 that reduced to 11%±6 after DIV10 (*Fig. 1D*). This result is consistent with the earlier reported trend(17).

**Figure 1.**
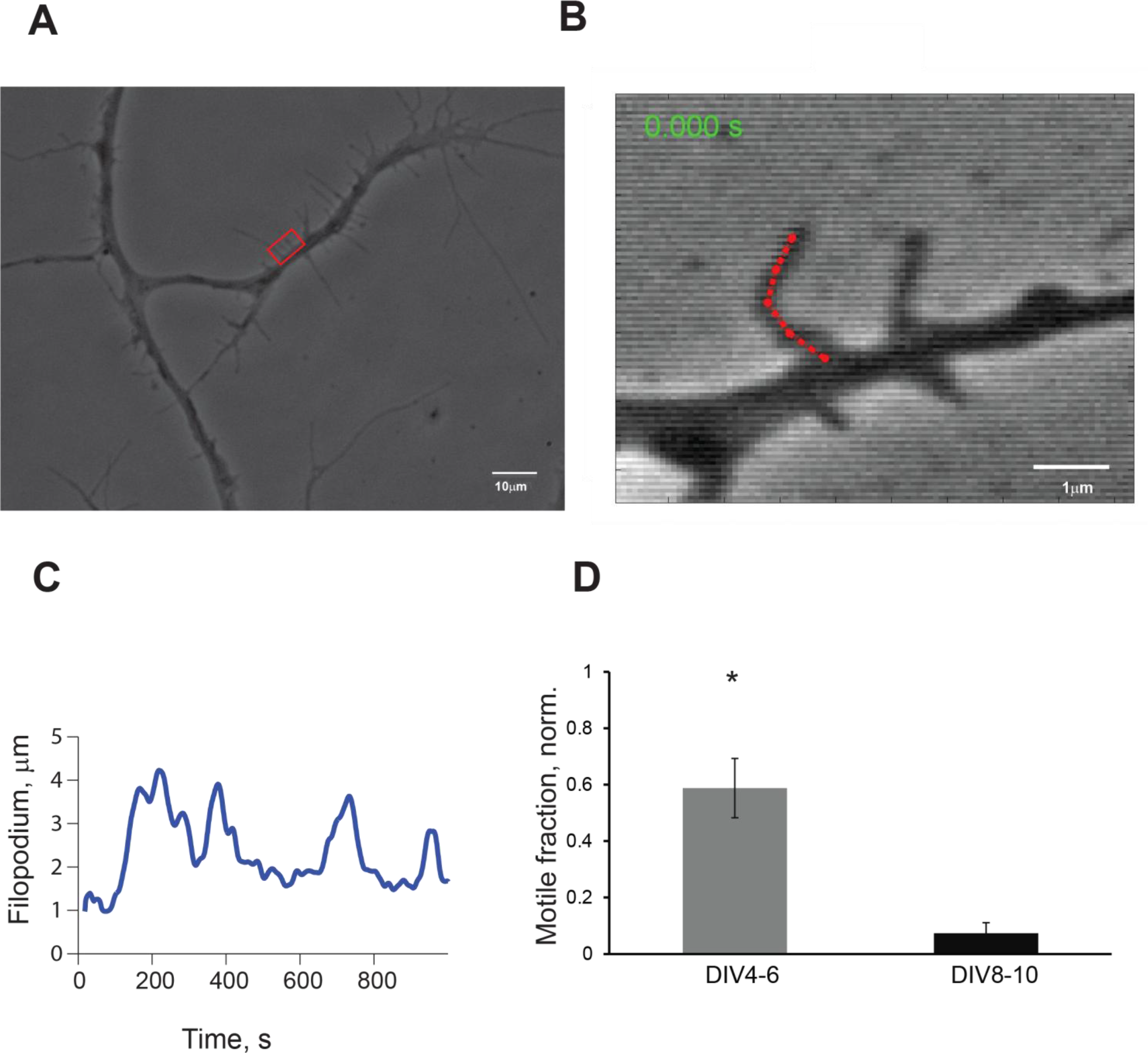
Analysis of filopodia motility with automated tracking software FiloTracker A. Phase-contrast image of hippocampal pyramidal neuron in culture, DIV5, with dendrites covered with filopodia; scale bar 10μm. B. High magnification image of boxed area shown in panel A during length tracking process in FiloTracker; red dotted line is a length estimate for the current frame; scale bar 1 μm. C. Output sample of length measurements from the filopodia on B (red), acquisition rate= 1f/s, periodicity of length is ~ 122±19s. D. Comparison of motile fraction of all filopodia in imaging session (t=30 mins) at DIV4-9 and DIV8-10 (t-test P<0.05).

In all filopodia trajectories with continual dynamics, we calculated the protrusion rates (length increase per time during extension) and retraction rates (length decrease per time during retraction). We determined that the lengths fluctuate with similar protrusion and retraction rates of 0.37±0.30μ/s and 0.32±0.24μ/s in DIV4-7, respectively.

We hypothesized that feedback among the actin retrograde flow (ARF), myosin activity and substrate adhesion are the principal factors that drive the range of filopodia behaviors observed in primary neuron culture, and a mathematical model was developed to quantitatively explore this hypothesis.

## Formulation of a model based on experimental constraints

We considered a minimal number of biophysical mechanisms as sufficient to describe filopodium dynamics, with its extension driven by actin polymerization and retraction by myosin contraction. The total contractile force exerted within the filopodium is proportional to the total amount of bound myosin. When the filopodium length is short, there is only a small amount of bound myosin, and consequently a small contractile force. Therefore, polymerization of the actin at the tip causes the filopodium to extend. As the filopodium gets longer, the contractile force gets larger due to binding of myosin, and the tip extension slows. At some point, myosin contraction can outstrip polymerization and the filopodium shrinks. As shrinking proceeds, the density of bound myosin increases and accelerates contraction. Ultimately, myosin dissociation reduces the contractile force and the filopodium length reaches its minimum before extension resumes. It is therefore possible that standard actomyosin dynamics can account for all ranges of filopodia dynamics. However, other factors such as the viscoelasticity of the actin network and the adhesion of the filopodium to the substrate may affect the overall system behavior. These and other features of the system can be quantitatively explored with a mathematical model based on five essential processes (*Fig. 2*): a) polymerization of F-actin at the tip of the filopodium; b) binding and unbinding of myosin to F-actin; c) isotropic contractile stresses exerted by myosin on F-actin; d) viscous flow of F-actin (ARF) induced by these contractile stresses; e) friction between the filopodium and the substrate due to adhesion (Bausch et al., 1998).

**Figure 2.**
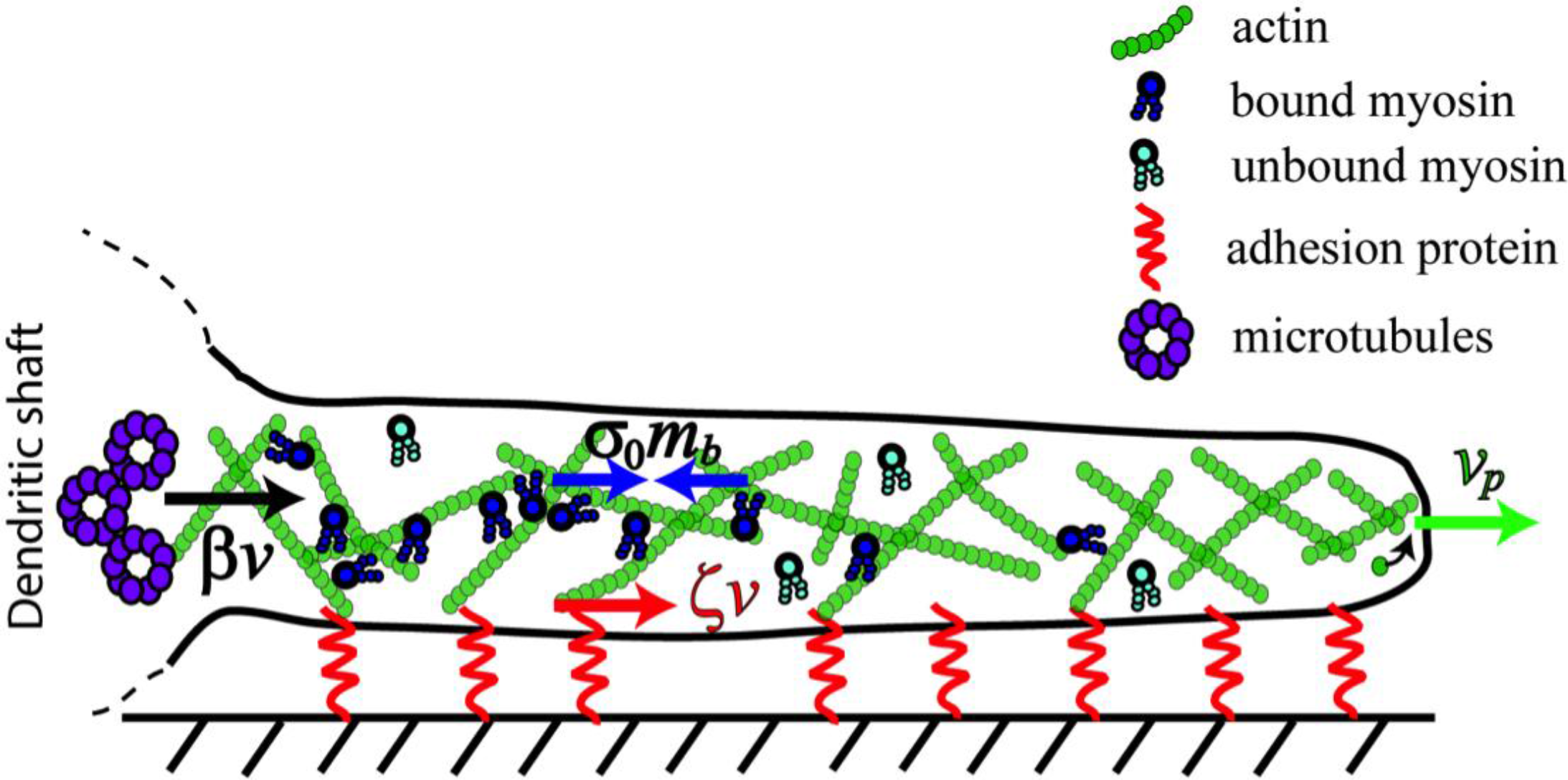
Cartoon that illustrates the balance of forces in a dendritic filopodium described by the minimal model. The model captures five essential processes: a) polymerization of F-actin at the tip of the filopodium; b) binding and unbinding of myosin to F-actin; c) isotropic contractile stresses exerted by bound myosin on F-actin; d) viscous flow of F-actin (ARF) induced by these contractile stresses and membrane tension; e) friction between the filopodium and the substrate due to adhesion. Bound myosin contractile stress is shown by blue arrows *σ_0_*, *m_b_*, *v_p_*-polymerization rate, *ζv*-the substrate adhesion force is denoted by the red arrow, and black arrow is *βv*-resistance force at the base due to microtubule network inside the dendrite. Unbound myosin freely diffuses inside the filopodium.

We implemented the model filopodium as a linear 1D object whose length *L*(*t*) varies with time. The state of the system is described by two variables, the distribution of bound myosin, *m_b_*(*x,t*) and the local velocity of the actin network *v*(*x,t*), *x ∈* [*0,L*], where *x*=0 corresponds to the base of the filopodium and x=*L* to the tip. Myosin can bind and unbind from the actin cytoskeleton with rate constants *k_on_* and *k_off_*, respectively. The unbound myosin is assumed to diffuse rapidly on the length scale of the filopodium and, on the timescale of filopodial dynamics, to be in equilibrium with the large reservoir contained in the adjacent dendrite; therefore its concentration, *m*, is assumed to be constant.

Bound myosin moves with the F-actin at the velocity *v*(*x*) and can also diffuse due to random motion of the actin filaments (21). The dynamics of the bound myosin is then described by the following continuity (transport) equation:

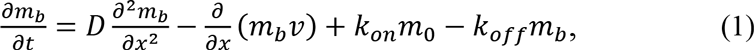
where *D* is the diffusion coefficient.

The bound myosin exerts a contractile stress on the actin network. We assume that this contractile stress is isotropic and proportional to the concentration of bound myosin. On the timescales relevant to filopodial motility, the actin network can be approximated as a viscous fluid with viscosity *η* (Bausch et al., 1998). Adhesion between the filopodial membrane and the extra-cellular matrix resists the movement of the filopodium. We account for this as a resistive force proportional to the velocity. Therefore, myosin contractile stresses will induce a viscous flow of F-actin governed by the mechanical balance equation,

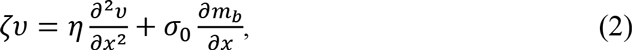
which takes into account the three major forces present in the system: viscous stress, the contractile stress exerted by bound myosin, *σ_0_m_b_*, and the resistive drag force, (*ζv*, due to adhesive interactions with the substrate. In *Eq.2*, *σ_0_* is a myosin contractility coefficient that should be approximately proportional to the maximum amount of work that can be done by a single myosin molecule.

*Eqs* (1, 2) are solved with the following initial conditions:

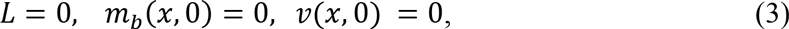
and *L*, *m_b_* and *v* allowed to evolve over time and along the one dimensional space. With regard to boundary conditions, we assume that F-actin and its bound myosin do not flow into or out of the dendritic shaft based on previous experimental findings (22) and possibly due to a barrier formed by microtubules in the adjacent dendritic shaft (this idea will be further explored below); therefore, the flux of bound myosin is zero at the base:

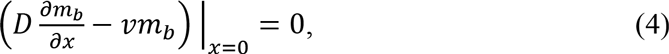

We use a resistive force proportional to the velocity, βν, to model the stress balance at the base of filopodium:

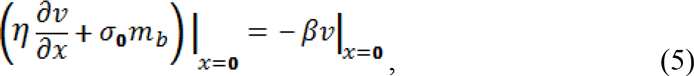

At the tip of the filopodium, membrane tension, *T*, exerts a force proportional to the curvature, *К*, onto the actin network:

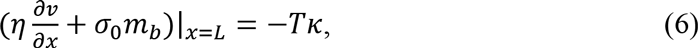
effectively, *TК* is a single parameter that is considered to have a negligible value.

The rate that the length of the filopodium changes is equal to the difference between the actin polymerization rate at the tip, *v_p_*, which is set to be independent of other system variables, and the retrograde velocity of the F-actin network. Thus,

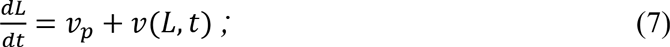

Since the actin at the tip is newly polymerized, myosin will not have bound yet, leading to the boundary condition *m*_b_(*L*,*t*) = 0. Model variables and parameters are listed in *Table S1*; the nominal parameter values shown in this Table are used throughout unless otherwise indicated.

## The model can recapitulate the experimentally observed motility behaviors

We analyzed equations (1–7) over a large range of parameters using a semi-implicit Crank-Nicolson scheme coded in MATLAB. We found that the model leads to three qualitatively different patterns of behavior: continuous increase of filopodium length, an equilibrium state with constant length, and periodic filopodium length fluctuations around a constant value. To begin, we focused on scenarios where the filopodium length fluctuated, similar to what has been observed during early neuron development (Ziv and Smith, 1996). To test the predictions of our model, we compared the model results to existing experimental measurements of actin retrograde flow (ARF) and myosin localization from live dendritic filopodia(Tatavarty et al., 2012). In these experiments, ARF was measured and averaged in motile and non-motile filopodia. We compared simulated ARF over protrusion and retraction phases and found striking agreement between the experiment and the model over a wide range of parameter values (*Fig. 3A*). Experimental data was used to set the parameters: *σ*_0_, *v_p_*, *k_on_*, and *k_off_* (*Table S1*). The values for unknown parameters: *ζ*, *η*, and *β* were obtained using a sensitivity analysis of the actin retrograde flow distribution *v(x)* with respect to each of these parameters in the vicinity of the “best fit” values (*Table S1*), such that the resulting ARF remained within the standard deviation of the experimentally-determined curve (*Fig. 3A*). The ranges for the unconstrained parameters that produce actin retrograde flow fit within the standard deviation bars are: *η*[34, 3000], *ζ*[118, 910], *β*[500, 50000]. Therefore, although we do not have experimental data to constrain the unknown parameters, the system is insensitive to the precise choice of these parameter values within the ranges indicated above. Interestingly, the parameter values that fit the experimentally measured ARF correspond to fluctuating and non-motile filopodia but do not constantly increase length.

**Figure 3.**
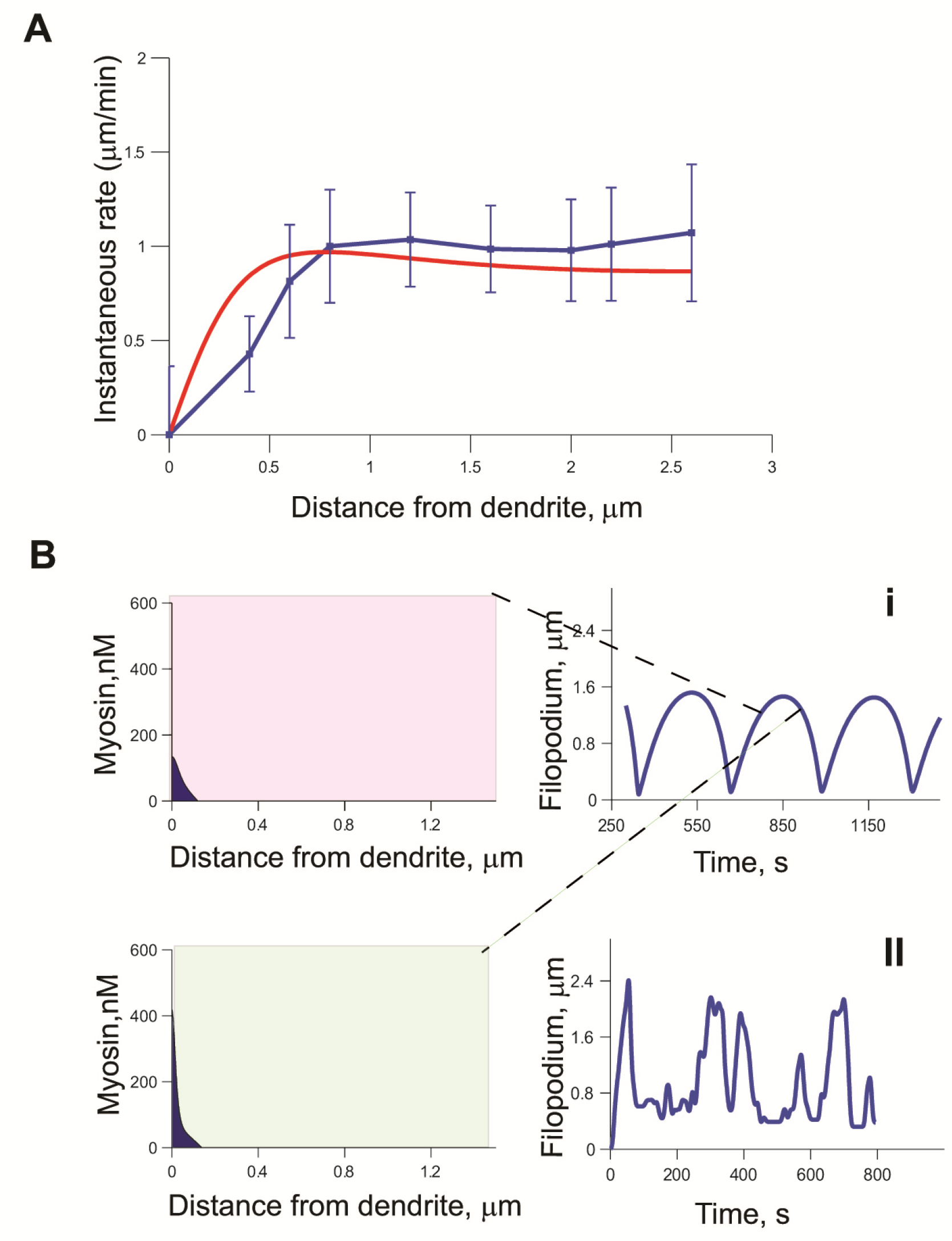
Model solutions correspond to experimental data. A. Fit of simulated actin flow velocity at steady state(red) (*Eq.1*) with experimental ARF distribution along normalized filopodium length (blue);(Tatavarty et al., 2012), mean± S.D. The confidence intervals for the unconstrained parameters that produce actin retrograde flow fit within the standard deviation bars are: η[5, 250], ς[0-120], β[500, 100000], σ_0_ =[1.1-1.8] L_0_ =1, k_off_=0.29, k_on_=0.27,v_p_=0.8, m_0_ = 200, η =100, ζ=100, σ_0_ =1.3, P=25000 B. i. Simulated filopodium length fluctuations with retrograde flow from A and corresponding myosin distribution taken close to the peak of the growth cycle (red insert) and during the retraction phase (green insert). ii) Tracking of live filopodium using automated software, aquisition rate = *1f/s*. L_0_ =1, k_off_=0.29, k_on_=0.27, v_p_=0.8, m_0_ = 200, η =100, Z=100, g_0_ =1.3, P=25000

Myosin activity is the main regulator of actin retrograde flow force. In the model, bound myosin is pushed by actin retrograde flow towards the dendrite, and accumulation of myosin at the base is largely due to the resistive barrier at the filopodium base, which prevents actin from flowing out of the filopodium (*Eq.5*).

Because of the barrier at the dendrite, the bound myosin remains highly localized at the filopodium base during all phases of the motility cycle. During the initial stage of protrusion, myosin binds to actin network which leads to generation of slow ARF and further promotes accumulation of myosin at the base of the filopodium. Finally, when the protrusion rate reaches zero at the peak of the growth phase, bound myosin localizes in high concentration right near the filopodium base (*Fig 3Bi*), while low concentrations are distributed along the filopodium length decaying to 0 at the tip. During the retraction phase the ARF is larger than during protrusion maintaining high concentrations of myosin at the filopodium base. Simulated actin retrograde flow profile that corresponds to the experimental findings gives rise to the filopodial length fluctuations (*Fig 3Bi*) comparable to the fluctuating filopodia tracked in culture (*Fig 3Bii*).

Thus, our model emphasizes that maintaining myosin localization at the base is required for persistent filopodial dynamics. To validate the model’s prediction, we compared the simulated myosin gradient with the experimental results in the active myosin distribution in live filopodia. Neurons were transfected with Eos-MLC ( myosin light chain) DNA construct(Tatavarty et al., 2012). Single-molecule PALM imaging showed localization of active myosin II near the base of the dendritic filopodia (*Fig 4A*). The probability of myosin light chain (MLC) residence at the base changed from almost 80% at DIV8-10 to 50% at DIV14-15 (*Fig. 4B*). Myosin localization was measured in dynamic and non-motile filopodia with both measurements showing maximum myosin concentration at the base (*Fig. 4B*). The averaged myosin distribution over the course of simulation (*Fig. 4C*) displays a base localization pattern similar to the one from the experimental results.

**Figure 4.**
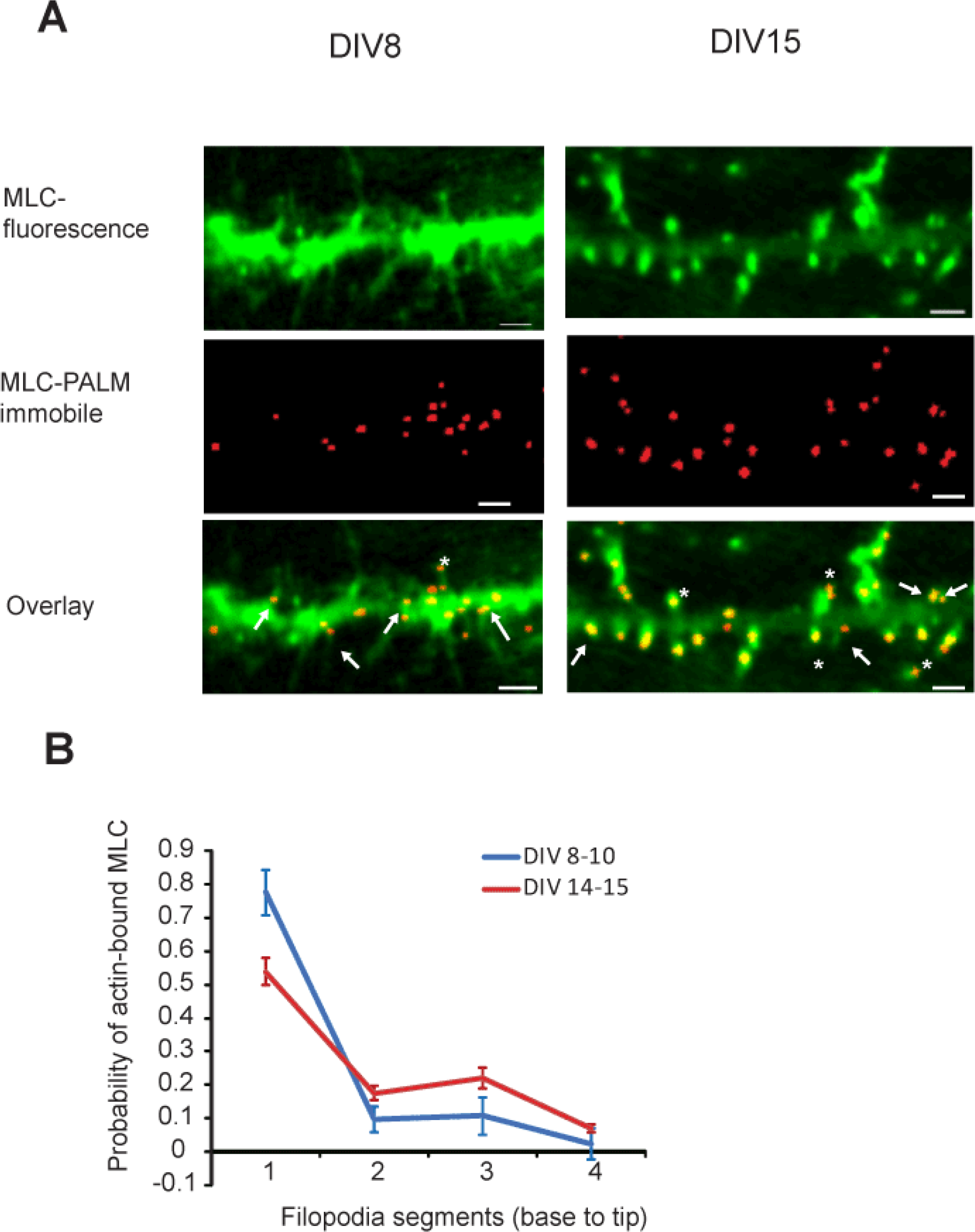
Localization of myosin (MLC) in dendrites using PALM imaging. A. Top, epifluorescence images of neurons transfected with Eos-MLC. Middle, corresponding PALM images constructed with only the immobile subpopulation of the molecules. Bottom,overlay of the top and middle images. Arrowheads indicate immobile MLC localization in the base of the filopodia. Asterisks indicate immobile MLC localization in the spine heads. Scale bar, μm. B. Probability of MLC to be found in one of the four segments of the filopodium during early development (DIV8-10, 26 filopodia) vs late development (DIV 14-15, 25 filopodia), error bars represent SEM; numbers indicate relative distance from the base. C. Simulation of distribution of bound myosin in filopodium averaged over all time points during oscillations shows accumulation of myosin at the base of the filopodium throughout simulation. Parameter values for the model output in A and B are: L_0_ =1, k_of_=0.29, k_on_=0.27, v_p_=0.8, m_0_ = 200, η =100, Z=100, σ_0_ =1.3, P=25000

While the filopodium is filled with branched filamentous actin, the dendrite contains a prominent microtubule network, as shown by electron microscopy images (*Fig 5A*). The dense network of microtubules runs uninterrupted in the dendrites orthogonal to the filopodial actin filaments forming a barrier. We hypothesize that the microtubules at the base of the filopodium function as a barrier to retain bound myosin and actin inside the filopodium (*Fig. 5B*), as expressed in the boundary condition of our model (*Eq. 4, 5*). The noflux condition for the actomyosin network at the filopodium base (*Eq. 4*) prevents filopodium collapse at the end of the retraction cycle, and is required for filopodium stability. By Newton’s law, the microtubule network has to exert some stress on the actin network proportional to the retrograde flow generated in the filopodium. However, it is unclear, if the stress from the microtubule network in the dendritic shaft plays any role in filopodia motility. Thus, to test the effect of the resistance force at the filopodium base we varied the resistance parameter *β* and observed its effect on filopodia dynamics. Values of *β* > 500 resulted in motile filopodium with regular length oscillations (*Fig. 5Bi*), while removing the stress at the base by setting *β* = 0 produced non-motile filopodia (*Fig. 5Cii*). Therefore, large *β* values dramatically increased the range of other unconstrained parameters that result in oscillations (*Supplemental materials*, *Fig.S1*). Interestingly, the steady state filopodial length for large and small *β* values remained the same, which is consistent with the no-flux assumption of the boundary condition (*Eq. 4*). Furthermore, treating the filopodia with small doses of nocodazole, a microtubule dissolving drug, abolished filopodia motility and significantly reduced their number (*Supplemental materials, Fig.S2*). In conclusion, our data and model are consistent with the hypothesis that the force at the base of the filopodium, proportional to the actin retrograde flow in the filopodium, stabilizes filopodial length and promotes filopodial motility.

**Figure 5.**
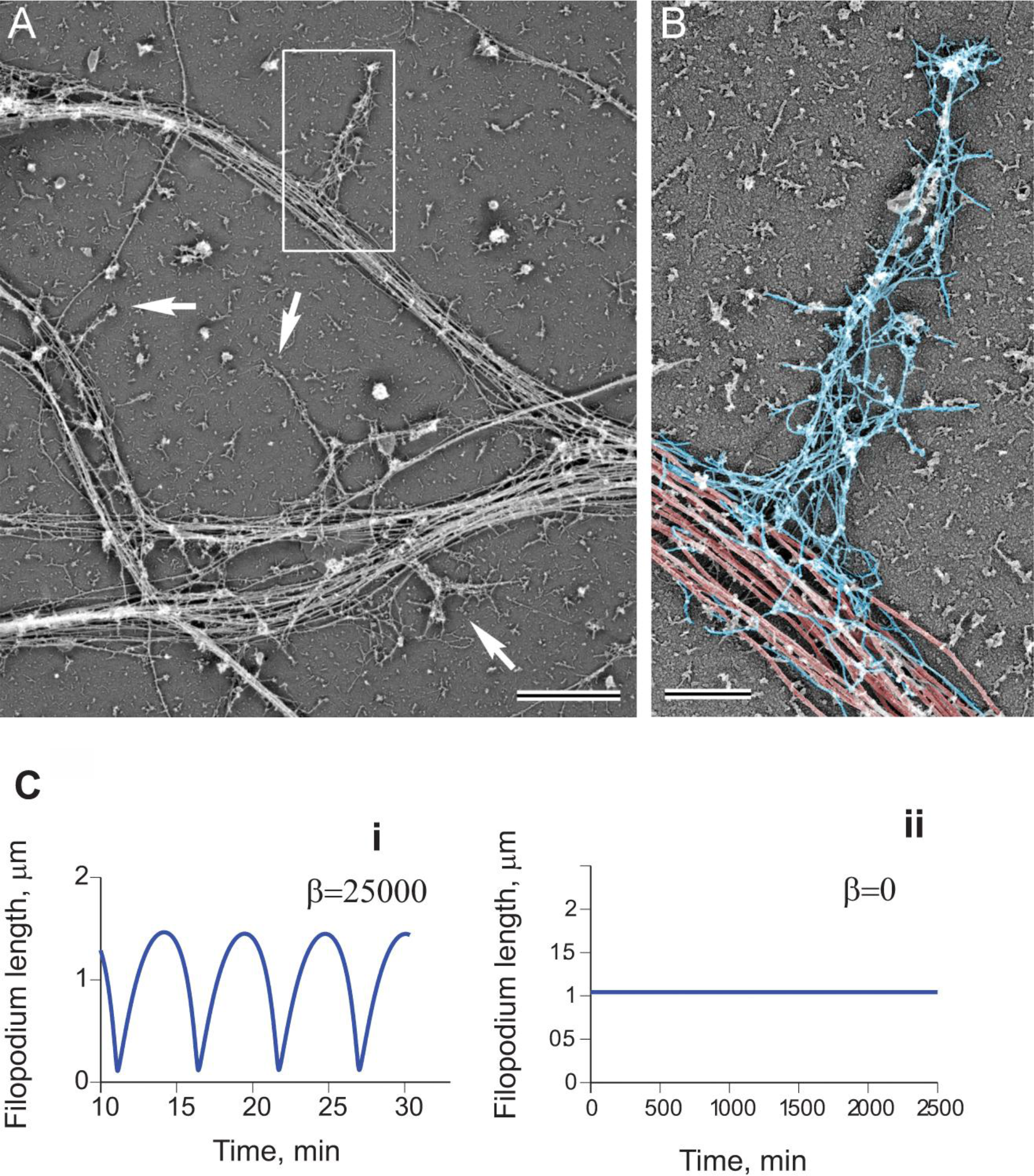
Filopodia dynamics depends on the resistive force applied at the base by the microtubule network. Cytoskeletal organization of dendritic filopodia revealed by platinum replica electron microscopy. (A) A network of dendrites from hippocampal neurons cultured for 8 days in vitro. Dendritic filopodia (boxed region and arrows) reside on dense arrays of microtubules in dendrites. (B) Dendritic filopodium from the boxed region in A is color-coded to show microtubules in the dendrite (red) and actin filaments in the filopodium (blue). Scale bars: 2 um (A) and 0.5 um (B). C. Simulated filopodia dynamics with i) and without ii) resistive force at the barrier. i) koff=0.29, kon=0.27,vp=0.8, m0 = 200, η =100, σ_0_ =2.5, P=25000 ii) koff=0.29, kon=0.27,vp=0.8, m0 = 200, η =100, σ_0_ =2.5, P=0

In summary, simulated actin retrograde flow and myosin dynamics recapitulated dynamic behavior of dendritic filopodia observed in live imaging experiments. We next explored whether the model can reproduce the effects of experimental manipulations. We have chosen a series of experimental manipulations that have direct correlates to the key parameters of the model.

## Actin polymerization rate regulates filopodia motility

Cytochalasin D (CD) is a pharmacological agent that disrupts actin polymerization by blocking G-actin incorporation into existing filaments. To understand the effect of polymerization rate on filopodia motility, we designed experiments to systematically decrease the actin polymerization rate with CD. We then compared these experimental results with results from our simulations where we decreased the polymerization rate *v_p_*.

The application of 20 nM CD abolished filopodia motility at DIV9 (*Movie S2A*). Treatment of neurons with lower concentrations of CD resulted in dose-dependent attenuation of filopodia motility (*Movie S2B*). While 20nM cytochalasin D treatment “froze” all motile filopodia, a 5nM dose of CD resulted in a 20% reduction in the number of motile filopodia compared to control. (*Fig. 6A*). Smaller concentrations of cytochalasin D (<5nM) did not produce statistically significant effects on the motility of dendritic filopodia at DIV4-7. The low dose and short time over which cells were exposed to CD assured that the actin cytoskeleton remained intact during our measurements (*Fig. S4*).

**Figure 6.**
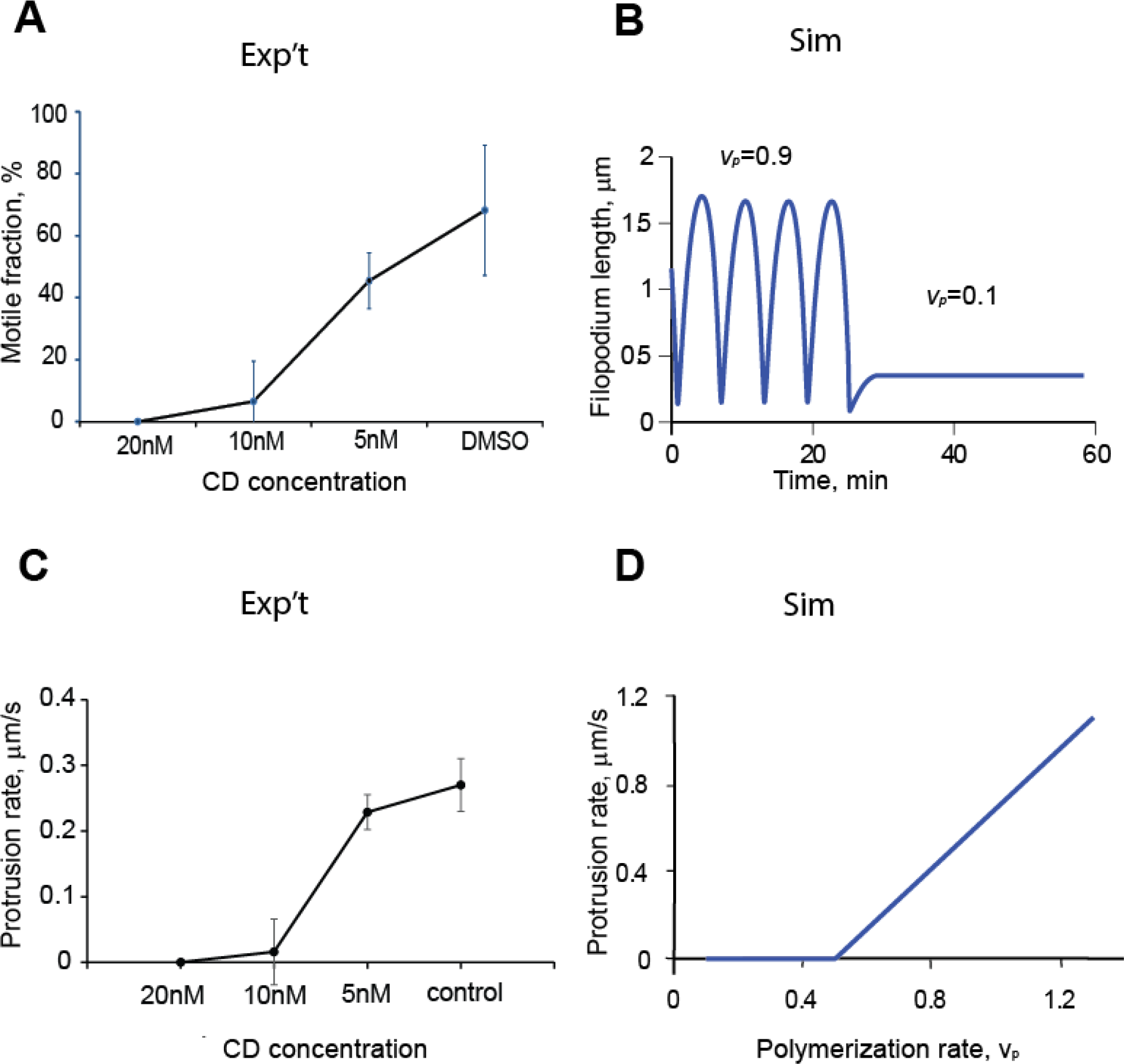
Effect of actin polymerization rate on filopodial motility. A. Percent motile filopodia as a function of CD concentration; one-way ANOVA P<0.001 for the 20nM dose compared to each of the lower doses (201 filopodia, 21 neurons, error bars are SD). B. Simulated changes in filopodial motility after decreasing actin polymerization rate, *v_p_* from 0.8 to 0.1. Other parameter values are: koff = 0.29, kon = 0.27, m_0_ = 200, η =100, Z=100, σ0 =1.3, P=25000 (*Eq.7*) C.Effect of CD treatment on the protrusion rate of motile filopodia, one-way ANOVA P<0.001, for the 20nM dose compared to each of the lower doses (124 filopodia, 18 neurons; error bars are SD). D. Protrusion rates from simulated filopodial dynamics using range of polymerization rate values, vp [0-0.8].․ Other parameters used in the simulations koff=0.29, kon=0.27,m0 = 200, η =100, ¶=70, σ0 =1.3, β=25000. The simulations were initiated with bound myosin (m_b_) = 0 and Length (L_0_) =1μm.

Since cytochalasin blocks incorporation of actin monomers into F-actin, the overall motility inhibiting effect of cytochalasin is not surprising (*Fig. 6A*). However, we specifically expected, that the decrease in protrusion rate is reproduced in the model simulations by perturbing polymerization rate *v_p_*. A step-wise reduction of the parameter *v_p_* from 0.8 to 0.1 resulted in essentially non-motile filopodia with the steady-state lengths four-fold smaller than the average lengths of filopodia at *v_p_*=0.8 (*Fig. 6B*). The Experimental data showed a gradual decrease of protrusion rates in dendritic filopodia treated with increasing CD doses (*Fig. 6C*). The trend we observed is reproduced by our mathematical model (*Fig. 6D*).

## The effect of substrate adhesion on filopodia dynamics is biphasic

The drag force, generated by substrate adhesion, inhibits actin retrograde flow and supports polymerizing actin filaments in an elongating filopodium (23). Poly-L-lysine is a positively charged polymer that effectively mediates adhesion in cultured neurons. PLL stimulates integrin-substrate binding via syndecans (24), and a range of pLL concentrations is used to obtain viable neurons in culture.

We plated hippocampal neurons on glass coverslip dishes coated with pLL concentrations of 0.01, 0.05, 0.5 1.0, or 10 mg/ml and measured filopodial density, and protrusion and retraction rates using our custom written tracking method described in *Fig.1*. The highest number of motile filopodia was measured at conc. 0.05mg/ml (*Fig. 7A*), with significantly smaller values at conc. 0.01mg/ml and 1.0 mg/ml (*Movies S3A-C*). Because protrusion and retraction rates were the same for the filopodia grown on corresponding pLL substrate, all rates were compared together (*Fig.7B*). Motile filopodia grown on pLL applied at conc. 0.05mg/ml had the highest protrusion and retraction rates of 0.37±0.080 μm/min and 0.32+0.054μm/min respectively, while filopodia grown at the other concentrations showed a significant reduction in protrusion and retraction rates (*Fig.7B*).

**Figure 7.**
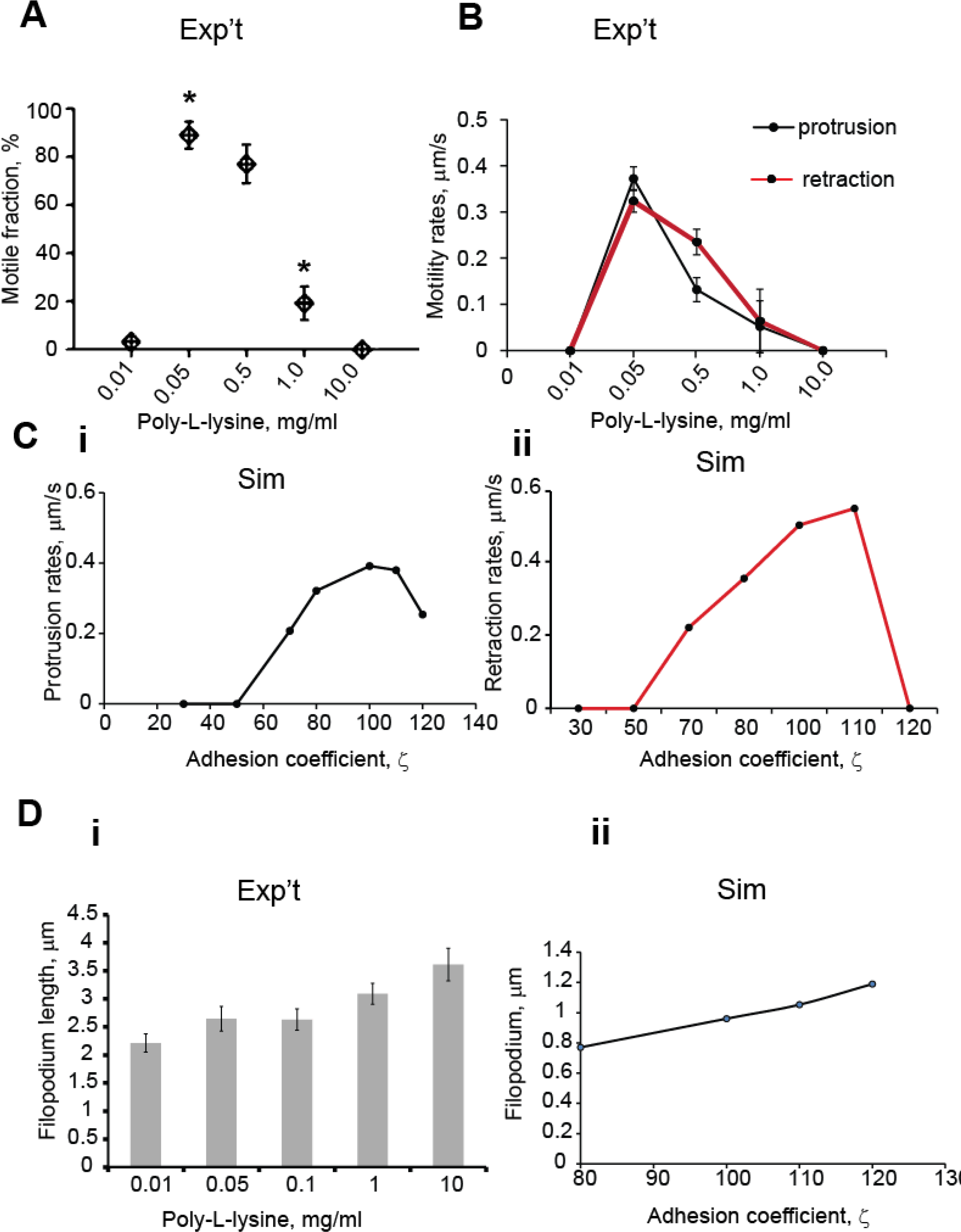
pLL-substrate adhesion strength regulates filopodial motility. A. The highest number of motile filopodia was observed on substrates with medium pLL concentration; using one-way ANOVA, motile fraction at 0.05 and 5 mg/ml were not significantly different from each other, but they were each significantly different than the filopodia at 0.01, 1.0 and 10.0mg/ml (433 filopodia, 36 neurons, P<0.001; ANOVA, error bars are SEM). B. Protrusion and retraction rates from filopodia grown on substrates of different pLL concentration (DIV4-5, 316 filopodia, 20 neurons); using one-way ANOVA, motility rates at 0.05 and 0.5 mg/ml were not significantly different from each other, but they were each significantly different than the less motile filopodia at 0.01, 1.0 and 10.0mg/ml (P<0.001; error bars are SD). C. Protrusion and retraction rates from simulated filopodia within adhesion coefficient range (30:180) demonstrate bimodal relationship between adhesion forces and filopodial dynamics. The simulations are initiated with bound myosin (m_b_) = 0 and Length (L_0_) =1μm; they both quickly approach non-motile steady state lengths, bracketing the intermediate ζ values that produce oscillations. Other parameters in these simulations are: koff=0.29, kon=0.27,vp=0.8, m0 = 200, η =100, σ_0_ =1.3, P=25000. D. Comparison of filopodial length change in modeling and experimental results: i) average filopodial lengths on the range of pLL concentrations (433 filopodia, 36 neurons, ANOVA, P<0.005; error bars are SEM; Tukey post hoc test: Group 0.1mg/ml has mean not statistically different from other groups; all four groups have means statistically different from each other t). ii) simulated filopodia lengths in the range of the drag force parameter ζ[10,180]

To examine whether these results correlate qualitatively with the modeling results, we simulated filopodia dynamics over a range of the drag force parameters *ζ*. A greater traction force with increasing drag force parameter resulted in changes in actin retrograde flow across the adhesion drag force range. As a result of this, the simulated protrusion and retraction rates in filopodia followed a bimodal trend corresponding to the net sum of forces

At *ζ* values below 30, the force due to friction was insufficient to generate motility and resulted in non-motile short filopodia, consistent with observations on live filopodia grown on low pLL concentrations. The protrusion and retraction rates were decreased compared to nominal case (This phenotype can be explained by lack of traction force and inability to generate sufficient actin retrograde flow necessary for retraction. Similarly, filopodia plated on poly-L-lysine taken at low concentrations were not able to undergo regular motility cycles and flailed during protrusion and retraction.

Filopodial protrusion rates decreased at *ζ*>110, and no regular motility was observed (*Fig. 7Ci*). Interestingly, the retraction rates were greater compared to protrusion rates (*Fig. 7Cii*). At larger values of the drag coefficient, friction forces counteract actin retrograde flow, and the resulting protrusion rate is small and insufficient to maintain retraction phase which halts motile cycle. This result is congruent with the observation that filopodia grown on poly-L-lysine taken at high concentrations were straight and immobile.

Since the effect of drag force coefficient on the filopodial motility is quite dramatic, we expected that filopodial lengths would also be affected by the changes in substrate adhesion. Indeed, when comparing experimental measurements between the lowest and highest PLL concentrations, filopodial lengths increased almost twice (*Fig7.Di*).

The increasing length trend was reproduced in simulations across the range drag force parameter *ζ* [0:200]. At *ζ*>109, the steady-state length of non-motile filopodia was greater than the average lengths of motile filopodia at lower values of *ζ*(*Fig. 7Dii*).

Thus, in simulated filopodia as well as in live neurons, intermediate values of adhesion drug coefficient produced filopodia with the greatest protrusion/retraction rates undergoing regular cycles of protrusion and retraction.

## Dendritic Filopodia Lengths and Motility are Increased by Blebbistatin Treatment

Experimentally, it has been shown that non-muscle myosin is a key regulating force in filopodia dynamics, and myosin inactivation by pharmacological agents, such as blebbistatin, abolishes filopodia motility (12, 25). The force opposing protrusion is generated by myosin contractile activity on the actin network and it affects both protrusion and retraction rates of the motile system. To evaluate how the myosin-actin interaction proposed in the model predicts and explains the live filopodia motility, we treated dendritic filopodia with a range of blebbistatin concentrations. To minimize the cytotoxicity and nonspecific effects of blebbistatin(26), we limited the upper concentration to 2.5μM (*Movie S4A*); at 10μM or higher, we observed collapse of filopodia and membrane blebbing.

The result of gradual myosin inactivation on filopodia dynamics was best captured by comparing filopodial lengths (*Fig. 8A, B*) and retraction/protrusion rates(*Fig. 8C, D, E, F*) in the experiments and simulations. This is because the protrusion/retraction rates, and length measurements (as opposed to the fraction of motile filopodia) are most directly compared to the output of our deterministic partial differential equations simulations. In the experiment, the filopodia were significantly longer in cells treated with 2.5μM blebbistatin, when compared to control (*Fig. 8A*). This is due to the fact that blebbistatin treatments resulted in increased retraction (*Fig.8C*) and protrusion rates(*Fig.8D*), as compared to the control. To explain the observed trends in filopodial motility rates and lengths in terms of the drug effect on myosin activity, we looked at how simulated filopodia dynamics change when the myosin on-rate parameter, *kon*\**m0*, is decreased.

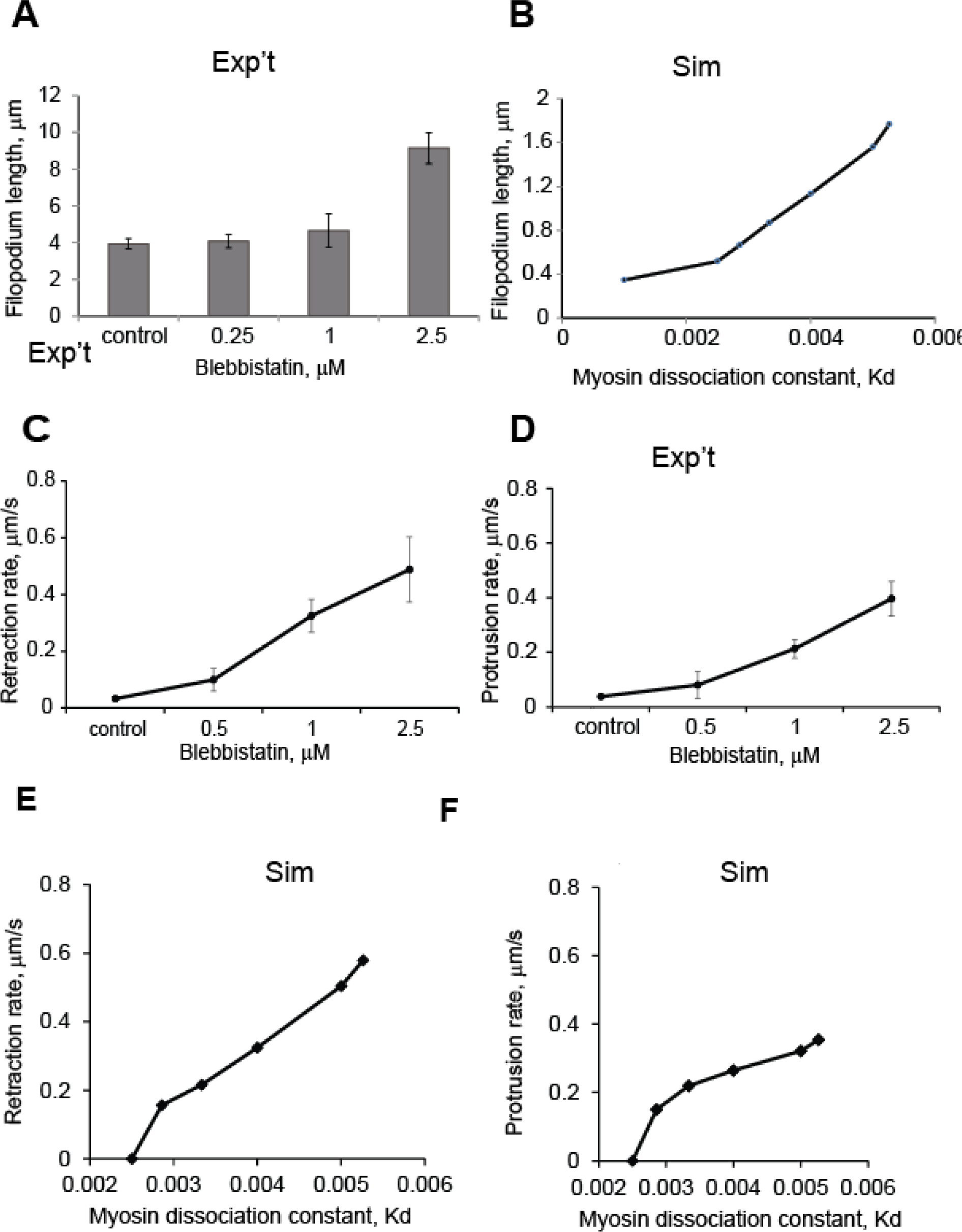

**Figure 8.**
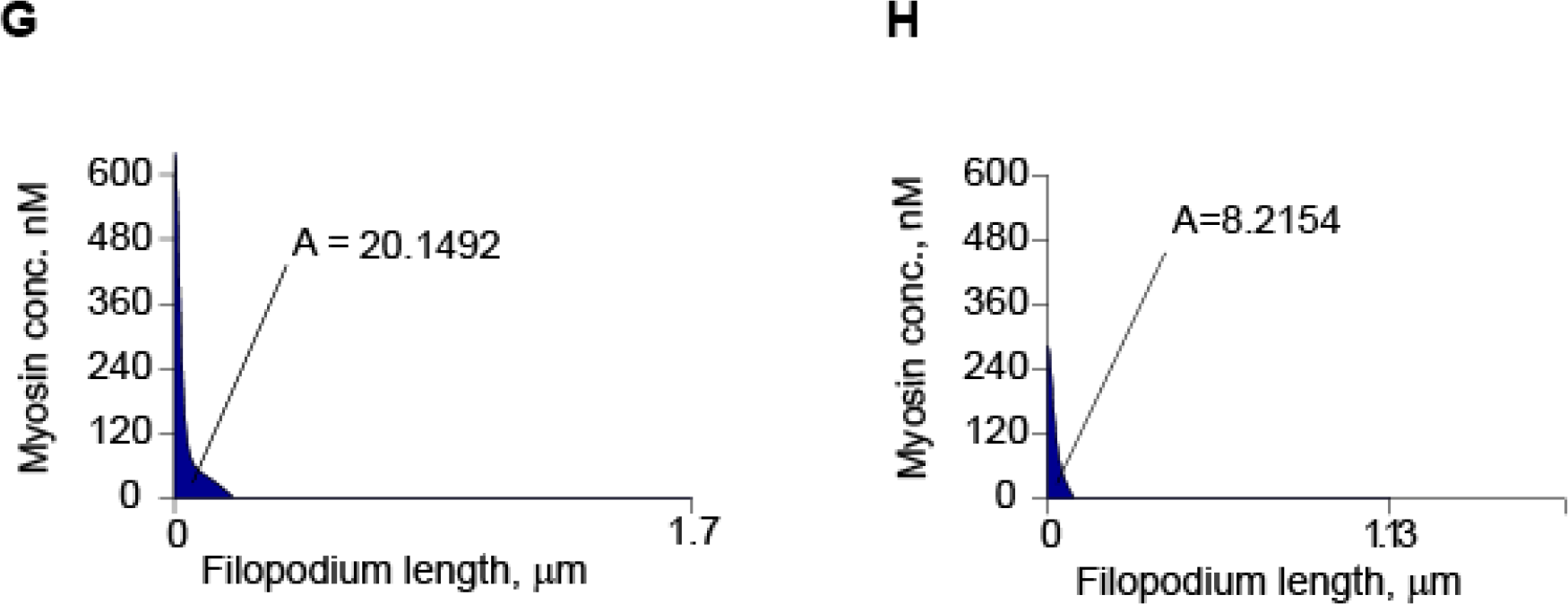
Dependence of filopodial lengths and retraction rates on myosin on-rate. A. Filopodial lengths after treatment with blebbistatin; significantly longer filopodia were observed after treatment with 1.0μM and 2.5μM doses of blebbistatin, when compared to control, one-way ANOVA P<0.0001 (95 filopodia, 12 neurons), error bars are SEM. B. Simulated filopodial lengths on the range of Kd [0.001,0.0055] C. Filopodial retraction rates after treatment with blebbistatin;the highest retraction rate was observed at blebbistatin treatment of 2.5μM (DIV4-5, 78 filopodia, 12 neurons, ANOVA, P<0.001. D) Filopodial protrusion rates after treatment with blebbistatin; the highest protrusion rate was observed at blebbistatin treatment of 2.5μM (DIV4-5, 78 filopodia, 12 neurons, ANOVA, P<0.001. E) Simulated filopodial retraction rates on the range of Kd [0.002,0.0055]; the highest retraction rate was observed at the highest Kd. Parameters used for the simulations are: k_off_=0.29, k_on_=0.27,v_p_=0.8,, η =100, Z=100, σ_0_ =1.3, m0=190-600, P=25000. F) Simulated filopodial protrusion rates on the range of Kd [0.002, 0.0055]; the highest protrusion rate was observed at the highest Kd. Parameters used for the simulations are: k_off_=0.29, k_on_=0.27,v_p_=0.8, · =100, Z=100, σ_0_ =1.3, m0=190-600 β=25000. G) Maximum myosin concentration at the peak of motility cycle (maximum length) at Kd = 0.005, A-total number of myosin molecules. H) Maximum myosin concentration at the peak of motility cycle (maximum length) at Kd = 0.0025, A-total number of myosin molecules.

Blebbistatin binds to myosin inADP-Pi state and interferes with the phosphate release process, locking myosin in the actin-unbound state (Kováks, et al., 2004). Thus, the effect of blebbistatin can be modeled as a decreased *m0* in *Eq. 1*, which effectively decreases the on-rate for formation of bound myosin, or, equivalently, increases the effective dissociation constant, Kd. In simulations, instantaneous protrusion rate was calculated at the half point between minimum and maximum lengths in the oscillation, and instantaneous retraction rate was calculated at the half point between the maximum and minimum lengths of the oscillation. In simulated filopodia dynamics, lower myosin on-rate resulted in longer filopodia compared to the control on-rate (*Fig.8B*). The protrusion and retraction rates in these filopodia were both increased compared to the nominal on-rate, while the period of the simulations was significantly greater. The time it took for the myosin to accumulate at sufficient amounts to induce retraction increases with on-rate reduction (270s with K_d_=0.004 and 427s with K_d_= 0.005), as would be expected; however because of the increased duration of the protrusion phase, there is actually sufficient time for a significantly increased level of bound myosin at the peak of an oscillation (*Fig.8G, H*), resulting in a faster retraction phase. In other words, the longer duration of the growth phase as well
as the longer length of the filopodia allow more myosin to accumulate despite the higher Kd, resulting in faster retraction (*Fig.8E*). Of course, if we were to decrease Kd for myosin binding still further, the system would stop oscillating and protrusion would proceed unchecked (not shown).

## Discussion

In this study, we numerically simulated and experimentally characterized two types of dendritic filopodia, non-motile and fluctuating. We demonstrated that a complex interplay among the actin retrograde flow, myosin contractility and substrate adhesion regulate filopodia dynamics observed in primary neuron culture. Consequently, we formulated a simple mechanism that explains filopodia transition from motile to immotile state. Understanding filopodia dynamics is a necessary step for developing insight into the dendritic spine plasticity and stability. Analyzing the filopodia motility in high-resolution is of great utility for identification of cellular phenotypes in neurodevelopmental disorders and development of drug-screening platforms.

We found that filopodia protrusion rate is highly sensitive to actin polymerization. Cytochalasin inhibits polymerization by blocking free barbed ends resulting in reduced filopodia protrusion rate. Similarly, in neuronal growth cones, 5μM cytochalasin B (CB) treatment resulted in reversibly blocked motility (Medeiros et al., 2006). The drug produced a dose-dependent reduction in filopodia protrusion and retraction rates, and steady state filopodia length. Thus, actin polymerization rate at the filopodium tip is one of the key parameters that determines the filopodia dynamics and lifetime. Continuous depolymerization rate has been reported at the dendritic filopodia base, where accumulation of myosin takes place, and it is significantly smaller than the polymerization rate (Tatavarty et al., 2012). While we did not explicitly include depolymerization rate in the model, we assumed that the v_p_ parameter at the filopodia tip is a net actin polymerization rate and combines the polymerization and depolymerization rates. Because only the polymerization rate at the leading edge is inhibited by CD, the depolymerization rate is not affected and contributes to the filopodia retraction rate. Therefore, the model can reproduce the experimental results very well with the simplifying assumptions about the rates.

Next, we showed that traction force due to substrate adhesion promotes filopodia motility on surfaces with intermediate adhesiveness. Filopodia protrusion/retraction rates and motile fraction exhibited a biphasic dependence on adhesion strength, with filopodia at intermediate adhesion strength displaying the highest motility rates. The results of simulations also suggested that protrusion and retraction rates in dendritic filopodia are highly sensitive to the strength of substrate adhesion. At low and high values of the adhesion coefficient the actin retrograde flow was weaker producing stable or slightly motile filopodia. The highest simulated motility rates were observed at intermediate values of traction force. These results are consistent with the trend in actin retrograde flow measurements on various adhesion strengths in axonal growth cones (Geraldo and Gordon-Weeks, 2009). Similarly, actomyosin-based keratocyte motility on fibronectin, also shows a bimodal distribution of locomotion speed on substrates of increasing adhesion (Barnhart et al., 2011). However, while the bimodal trend is reproduced by the model, the exact relationship between PLL concentration and adhesion coefficient value is not known.

Interestingly, the data suggests that the filopodia lengths are slightly greater at the high concentrations of PLL in the substrate. This effect can be due to the fact that while the retraction and protrusion rates are slower, the force required to overcome the friction of the substrate increases with the stickiness of the substrate, and thus the minimal length that filopodium can reach at the end of retraction phase increases, which consequently leads to longer filopodia on average.

The treatment of adhesion in our model assumes that the traction force arises from uniform adhesion between the filopodium and the substrate. A more elaborate description that includes elastic properties of integrin-substrate linkage is needed to model stochastic events like slipping and tethering, which are not included in our model. Filament breaking can further attenuate filopodia formation on highly adhesive substrates. Nevertheless, a basic description of traction force correctly recapitulates the bimodal trend observed in the experiments.

The next important parameter of filopodia dynamics is the effect of myosin activity. Experimentally, we show that bound myosin accumulates at the base (*Fig. 4A–B*), and also consistent with the model simulations where bound myosin is highly biased toward the filopodium base during oscillatory dynamics (Fig.4C). This pattern reflects the fact that the actin filaments at the base region are the oldest and therefore have the most time to bind myosin. The myosin localization pattern maintains the actin retrograde flow, which in turn causes cyclic length contractions in filopodia.

To further explore the role of myosin for filopodium motility, we disrupted myosin binding to actin with low doses of blebbistatin (Loudon et al., 2006). Treatments with a range of increasing blebbistatin concentrations led to dendritic filopodia with increased protrusion rates (*Fig. 8D*), which is expected because protrusion is a competition between actin polymerization at the tip and actomyosin-dependent contraction. Similarly, the average length of filopodia also increases with increased blebbistatin concentration (*Fig. 8A*). Both of these responses to inhibition of myosin binding are recapitulated by the model (*Fig. 8F* and *8B*, respectively). However, unexpectedly and remarkably, retraction rates also increased with low doses of blebbistatin (*Fig. 8C*). While this results seems counterintuitive, it is predicted by the model (*Fig. 8E*). In addition, analysis of the model suggests a reason for this unexpected behavior: the longer duration of the fluctuations required to reach the maximum length when myosin binding is inhibited also gives the myosin more time to accumulate in the region of the filopodium base. Comparing *Fig. 8G* and *8H* shows that the accumulation of actomyosin is actually greater at the peak filopodial length when myosin binding is inhibited. Another way of thinking about this is to consider the oscillations as arising from a kinetic overshoot in the approach toward balance between actin polymerization and contractility; the overshoot is more severe when myosin binding is inhibited.

It is important to consider potential off-target effects of blebbistatin. While it has been reported previously that myosin inhibition with blebbistatin decreases myosin dependent F-actin disassembly and thus inhibits overall actin turnover rate (Ryu et al., 2006), it was also suggested that blebbistatin decreases actin polymerization rate as an off-target effect, which can cause the decreased motility in filopodia (Tatavarty et al., 2012). The findings also describe nonspecific effects by blebbistatin in *Dictyostelium* myoin II-knockout cells(Kolega, 2006). Indeed, we were unable to use blebbistatin concentrations in excess of 10μM in our experiments, as these concentrations caused complete loss of filopodia and even shrinkage of dendrites. To avoid any of these non-specific effects we limited our measurements to concentrations of 2.5μM and below. Therefore, we believe that effect of blebbistatin on myosin contractility in filopodia was highly specific.

Membrane tension has been shown to play a role in regulating cell motility in rapidly moving keratocytes (Lieber et al., 2013). While we did not directly probe the effects of membrane tension with our model, our results show that variations in membrane tension are not required to produce any of the dynamics reported here. The force from membrane tension is proportional to the curvature of the membrane. Therefore, this force can vary spatially across the membrane in a keratocyte However, for thin projections like filipodia, the tension force is set by the force required to pull out a membrane tether, which depends on the membrane surface tension and the bending rigidity of the membrane(Sheetz, 2001). Therefore, this force is expected to be relatively constant for dendritic filopodia and, thus, is not expected to regulate the dynamics of this system.

To summarize, our model reproduces experimental actin flow patterns, as well as the effects of varying myosin, adhesion and actin polymerization that lead to emergence of dynamic filopodia through the following mechanism. Consider a filopodium with a stable steady state length (*Fig. 9A*). In this scenario, the actin retrograde flow due to contraction and tension balance the polymerization rate. But this balance can only be attained if the myosin can enter and bind to the filopodial actin at a rate sufficient to keep up with the polymerization rate. Now consider a scenario where myosin binding and unbinding are slow compared to polymerization (*Fig.9B*). If the polymerization velocity is fast compared with the on rate of the myosin, then the filopodium will grow to be longer than the balance point that would have been achieved if myosin were fully bound to actin. As time progresses, though, the myosin will bind to a point where contractile velocity is equal to polymerization velocity. However, the delay caused by the slow myosin unbinding rate is such that the myosin continues to bind, leading to “overshoot” and producing a contractile stress that overcomes the polymerization velocity (*Fig.9B*). At this point, the filopodium will start to contract. As the filopodium contracts, myosin continues to bind, which causes the contractile velocity to increase. In addition, as the filopodium shortens, there is less external drag (because the total external drag is proportional to the length of the filopodium). Therefore, the contraction accelerates even further. In the absence of any other effects, the filopodium can end up shrinking to zero size. However, the viscosity of the actin prevents the filopodium from shrinking too quickly and this effect increases as the filopodium gets smaller. Therefore, the actin viscosity can prevent total collapse of the filopodium. Once the unbinding of myosin finally reduces contractility enough for polymerization to once again dominate, the filopodium begins to grow again and the cycle repeats.

**Figure 9.**
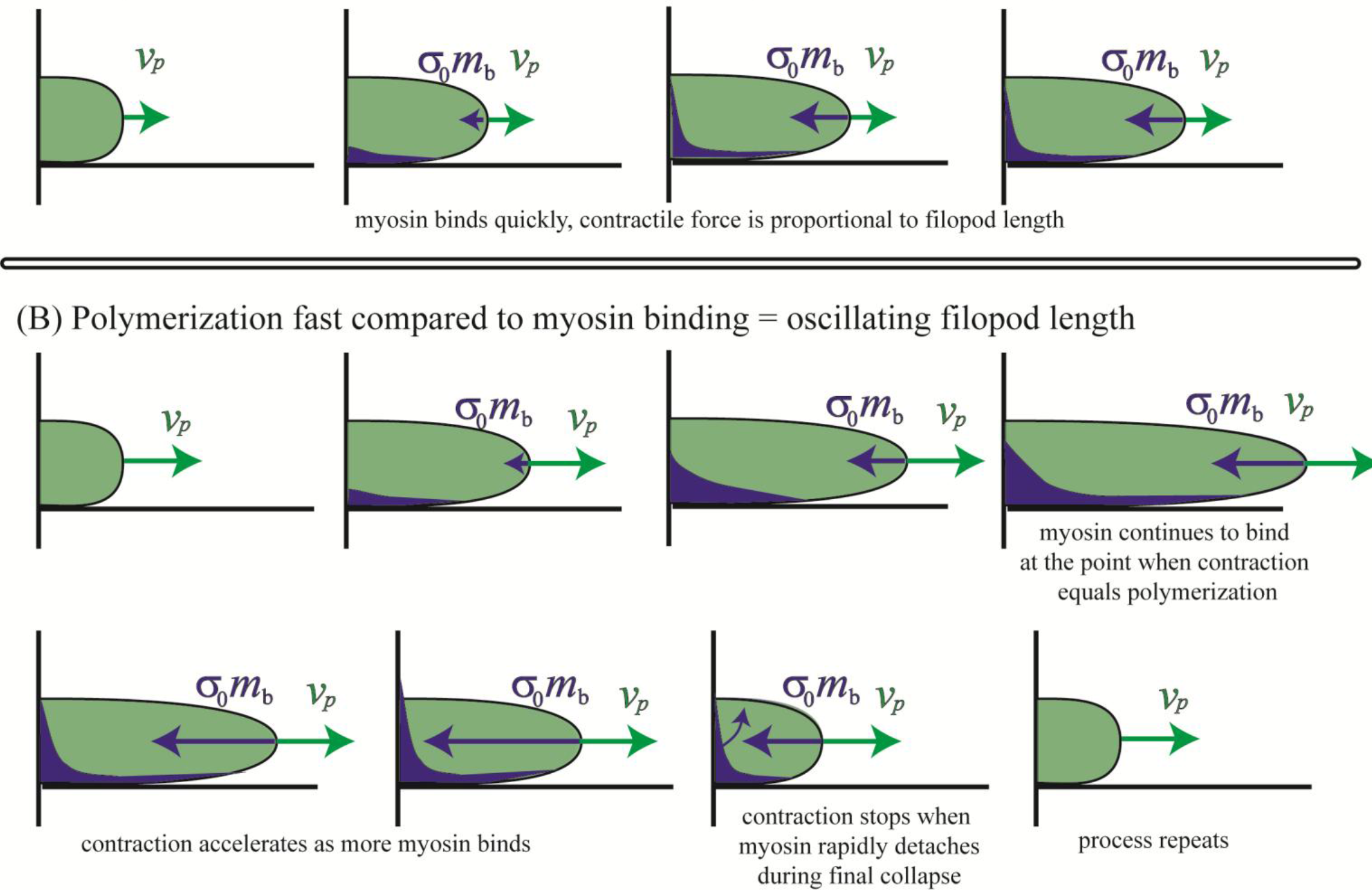
Summary of filopodium non-motile (A) and motile (B) behaviors.A. Non-motile filopodium. The length of filopodium does not changewhen myosin dynamics is fast enough to counteract the protrusion rate produced by actin polymerization. B. Motile filopodium undergoes protrusion-peak-retraction cycle. At small lengths the total amount of bound myosin is low; hence, retrograde flow is insufficient to overcome polymerization velocity, which results in gradual length increase. Filopodium reaches its maximum length when actin retrograde flow at the tip equals exactly to actin polymerization rate. Peak is followed by retraction stage, which is characterized by the large magnitudes of actin retrograde flow due to large amounts of bound myosin. The retraction lasts until the length is so small that the amount of bound myosin is low, which allows actin polymerization to increase the filopodial length and initiate the protrusion phase. This leads to the cyclical changes in filopodial length.

Whether filopodia dynamics and stability are important for spine formation is controversial (Portera-Cailliau et al., 2003, Hotulainen and Hoogenraad, 2010). Because the actin network inside the dendritic spine consists mainly of the same components as filopodia, we suggest that the factors that play important role in filopodia stability will be also important in spine morphogenesis and stability. Thus, in future studies we plan to focus on three central aspects of spine morphogenesis: metamorphosis of filopodia into spines, diversity of spine shapes and their stability. To gain insight into the regulation of spine shape, we will extend the 1D model to develop a 2D mechano-chemical model of spinogenesis, where the actin network is modeled as a viscoelastic fluid within moving boundaries. The 2D model will then be used to identify parameters and filopodia motility profiles that steer spinogenesis towards stable shapes.

## Materials and Methods

### Cell culture

Hippocampal neuron cultures were prepared from E18 rat (Sprague-Dawley) hippocampi obtained from BrainBits, Springfield, IL and plated following BrainBits Kit Protocol http://www.brainbitsllc.com/brainbits-kits-primary-neuron-protocol/. Briefly, cells were plated at 20-30,000 cells/well on MatTek (MatTek Corporation, Ashland, MA) glass bottom culture dishes that were thoroughly cleaned by sonicating sequentially in 10% HCl, 20% NaOH and Millipore water or in plasma cleaner and coated with poly-L-lysine (Sigma-Aldrich, St. Louis, MO) overnight before use. Cell cultures were maintained for up to 25 DIV in neurobasal medium supplemented with B27.Samples were imaged with Nikon Plan NA1.25x100 objective on DIV 3-12 on Nikon diaphot 300 inverted microscope. Images were collected with back-illuminated CCD camera (CH350, Roper Scientific) driven by Metamorph image acquisition and analysis software (Metamorph, Sunnyvale, CA).

### Drug treatments

For control conditions DMSO was added to Hibernate E, mixed well and the culture media was replaced with the DMSO solution at RT. The cells were imaged immediately following the media replacement for 20min. Blebbistatin(Sigma-Aldrich, St. Louis, MO) was used at 1.0, 2.5, 5.0, 12.5, 25, 50 μM final concentration in DMSO (2.5%) and Cytochalasin D (Sigma-Aldrich, St. Louis, MO) was used at 5.0, 10, 20 nM in DMSO (2%) final concentration at RT. At the start of the recording cell medium was replaced with the well-mixed solution of drug dissolved in DMSO and Hibernate E. The filopodia motility was recorded for 20 minutes after drug administration.

### Eos-MLC imaging

To express Eos-MLC fusion, a Gateway cassette (Invitrogen) was inserted to the C-terminal of the tdEos sequence (a gift from J.Wiedenmann, University of Ulm), which converted the original vector into a Gateway vector. An entry clone of the myosin regulatory light chain, Mlrc2, was purchased from Open Biosystems (Thermo Biosystems, Huntsville, AL). The Mlrc2 sequence was then subcloned into the tdEos Gateway vector using the LR clonase (Invitrogen) following the manufacturer’s procedure to produce the Eos-MLC fusion construct.

Single molecule imaging of Eos-MLC was performed using a modified epifluorescence microscope (Olympus IX81; Olympus, Tokyo, Japan) equipped with 60× microscope objective (numerical aperture, 1.45; Olympus) and a thermoelectric-cooled, electron-multiplying charge-coupled device camera (PhotonMax; Roper Scientific, Trenton, NJ). A 405-nm diode laser (Cube laser system; Coherent, Santa Clara, CA) was the light source for photoactivation. The MLC-green fluorescence of unactivated Eos was excited with the 488-nm laser line from an argon ion laser (CVI Melles Griot, Albuquerque, NM). Activated single molecules were imaged with a 532-nm, diode-pumped solid-state diode laser (Lambda Photometrics, Harpenden, UK). Time-lapse images were acquired every 0.3secs with 100-200ms exposure for each image. Image acquisition software was built on top of the μManager platform (http://micromanager.org). Image analysis was performed using a custom-built particle tracking algorithm as described before (Tatavarty et al., 2009). Only stationary single molecules of Eos-MLC (which moved less than 1 pixel) were considered for generating PALM images (MLC-PALM immobile), since this is the MLC-fraction which is bound to the actin filaments (Tatavarty et al, 2012). Filopodia with length 2μm or longer were chosen for analysis, and were binned into 4 segments, each of minimum 0.5μm. Individual immobile MLC molecule was assigned to a particular bin segment along the filopodia.

## Immunostaining

Cells were washed with PBS at RT and were fixed in gluteraldehyde 2% for 10min at 20°C, then washed with PBS for 30s and permealibized with Triton X-100 0.5% in PBS for 5min at 20°C. Cells are then rinsed with PBS twice at RT and incubated in BSA for 1hr. After 5min wash in PBS the cells were incubated with Acti-Stain 488 (Cytoskeleton Inc., Denver CO) or phalloidin 555 (Cytoskeleton Inc., Denver CO) for 30min.

Staining for β-tubulin was made as described in (33). Briefly, the cells were fixed with gluteraldehyde and permeabilized with Triton X-100. Imaging was performed on Zeiss LSM 780, a combined confocal/FCS/NLO system, mounted on an inverted Axio Observer Z1 with Plan Neofluar 63OilxNA1.25 objective.

## Platinum replica electron microscopy

Dissociated rat embryo hippocampal neurons isolated as described previously (Wilcox et al., 1994) were obtained from the MINS Neuron Culture Service Center (University of Pennsylvania, Philadelphia, PA). In brief, hippocampi were dissected from brains of Sprague-Dawley rat embryos at embryonic day 18-20 and dissociated into individual cells by incubating in a trypsin-containing solution. The cells were then washed and plated on poly-L-lysine-coated (1 mg/ml) glass coverslips at a concentration of 150,000 cells per 35-mm dish in 1.5 ml neurobasal medium (Gibco) with 2% B27 supplement. Sample preparation for platinum replica electron microscopy was performed as described previously (Svitkina, 2007; Svitkina, 2009). In brief, neuron cultures were detergent extracted for 20 sec at room temperature with 1% Triton X-100 in PEM buffer (100 mM Pipes-KOH, pH 6.9, 1 mM MgCl2, and 1 mM EGTA) containing 2% polyethelene glycol (molecular weight of 35,000), 2 μM phalloidin, and 10 μM taxol. Detergent-extracted samples were sequentially fixed with 2% glutaraldehyde in 0.1 M Na-cacodylate buffer (pH 7.3), 0.1% tannic acid, and 0.2% uranyl acetate; critical point dried; coated with platinum and carbon; and transferred onto electron microscopic grids for observation. Samples were analyzed using JEM 1011 transmission electron microscope (JEOL USA, Peabody, MA) operated at 100 kV. Images were captured by ORIUS 832.10W CCD camera (Gatan, Warrendale, PA) and presented in inverted contrast. Color labeling and image overlays were performed using Adobe Photoshop (Adobe Systems, Mountain View, CA), as described previously (Shutova et al., 2012).

## Filopodia length tracking

Custom-written software FiloTracker was used to track individual filopodia lengths. The use of an automated procedure excluded bias in selection of filopodia and increased the data throughput. Grayscale movies were segmented in ImageJ (National Institutes of Health, Bethesda, MD) and loaded into Matlab (MathWorks, Natick, MA). The individual filopodia length was measured and recorded into output file at each frame (dt=1s). Program output.mat and.csv files which were then imported into Matlab for further quantitative analysis. For description of the FiloTracker software, refer to the Supplemental Materials (Section 1). The tracks of filopodia motility were smoothed using Gaussian filter to remove noise and the protrusion retraction rates were computed using custom-written script.

## Computational analysis

The model is solved numerically using a semi-implicit Crank-Nicolson scheme coded in Matlab. Filopodium protrusion and retraction rates were computed using peakfinder function by Nathan Yoder, available on Matlab FileExchange: http://www.mathworks.com/matlabcentral/fileexchange/25500-peakfinder-x0–-sel–-thresh–-extrema–-includeendpoints–-interpolate-.

Filopodium length was estimated as maximum length from the motility track for each filopodium.

## Statistical analysis

Statistical analysis was conducted using one-way ANOVA tests with t-tests performed in Matlab. Error bars are mean ±SEM, unless noted otherwise.

## List of Supplemental Materials

1. **Supplemental Methods: Description of FiloTracker Software**
2. **Supplemental Figures and Tables** **Table S1. Parameter ranges resulting in ARF and myosin localization profile that match experimental data**
3. **Legends to Supplemental Movies**
4. **Supplemental References**

## Acknowledgements

We thank Dr. Boris Slepchenko, Dr. Yi Wu and Dr. Betty Eipper for valuable discussions on model validation, analysis and interpretations.

## Supplemental Materials

### 1. Supplemental methods. Description of FiloTracker Software

We developed an algorithm, implemented in Matlab, for tracking dendritic filopodial dynamics. The algorithm, which we call FiloTracker, can analyze a binary stack derived from DIC, phase contrast or fluorescence movies by automatically skeletonizing the filopodia into connected linear segments.

To produce a binary image for the algorithm two feature extraction steps were performed: edge-detection and foreground-background segmentation. At that point all the images in the stack are binary and ready for skeletonizing and tracking. FiloTracker then implements the annealed particle filtering algorithm originally developed to analyze articulated body motion in the computer vision field (Isard and Blake, 1998); the main idea is to compute the best configuration (the set of parameters that describes the object shape) as a statistical average over an ensemble of possible configurations, rather than searching for the best parameter set through some minimization procedure. Each filopodium in our method is represented by its skeleton, which is a nonself-intersecting polygonal chain with a fixed number, n, of segments of equal lengths. The skeleton is parametrized using the angles between the segments and the length of the segments. The default value of n is 4, but it may be adjusted by the user to achieve better results for extremely curvy filopodia. To initiate the skeletonization process, the user chooses filopodia in the first image of the stack by clicking pixel pairs at the base and tip of each filopodium. For each consecutive image, the algorithm first generates a sufficiently large random set of possible configurations in the vicinity of the skeleton from the previous frame; the only required constraint is that the base of all configurations is fixed. We then use pixel values from the new image to compute weights for these configurations, where weight is a measure of how closely a configuration lines up with the pixels in the new frame. Finally, in the new frame, the algorithm computes the configuration for the new skeleton as a weighted vector sum of all configurations from the original set. Full details on the mathematics behind the algorithm can be found in (Isard and Blake, 1998). Once the best fit filopodium is found on each consecutive frame, the length measurements are recorded into the output file. We found that image acquisition at 1 frame per second was sufficiently fast for highly precise tracking.

We validated our model with two independent test procedures. For the first, we generated a spatial simulation of a model filopodium that randomly grows, retracts and flails. The filopodium was simulated as a non-self-intersecting polygonal chain with a given bending modulus and given polymerization/depolymerization rates. Random force was applied at each node of the chain to generate realistic filopodia dynamics. Then, we used FiloTracker to measure the filopodium length. We compared the exact length values to lengths from the FiloTracker output and evaluated algorithm’s performance *(Fig.S5)*. In the second validation we tracked filopodia by hand and compared hand-tracked filopodia length measurements to FiloTracker output *(Fig.S6)*.

### 2. Supplemental Tables and Figures

**Table S1.** Parameter values for nominal case result in ARF and myosin localization profiles that match experimental data.

| Forces/Processes | Parameter | Physical significance | Value range | Units | Reference/source |
| --- | --- | --- | --- | --- | --- |
| Force due to viscous flow of actin | $\eta$ | Viscosity | 100 | $\frac{\text{g}}{\text{s } \mu\text{m}}$ | Sensitivity analysis |
| Myosin contractility | $\sigma_0$ | myosin contractility | 1.3 | $\frac{\text{pN}}{\text{mole}}$ | Norstrom et al. 2010, sensitivity analysis |
| Adhesion Force | $\zeta$ | drag force coefficient | 100 | $\frac{\text{g}}{\text{s } \mu\text{m}^3}$ | Sensitivity analysis |
| Myosin binding | $k_{\text{off}} / k_{\text{on}}$ | myosin binding rate | ~1 | n. a. | Norstrom et al. 2010 |
| Actin polymerization | $v_p$ | polymerization rate | 0.5-1.0 | $\text{s}^{-1}$ | Tatavarty et al.2012, |
| Initial filopodia length | $L_0$ | initial length | [0.01,1] | $\mu\text{m}$ | experimental data |
| Diffusion | $D$ | diffusion coefficient | 0.1 | $\mu\text{m}^2/\text{s}$ | Calculated from myosin random walk |
| Resistive force | $\beta$ | Resistive force coefficient | 25000 | $\frac{\text{g}}{\text{s } \mu\text{m}^3}$ | Sensitivity analysis |
| Unbound myosin concentration | $m_0$ | Unbound myosin | 200 | nM | Sensitivity analysis |
| Membrane tension | $T\kappa$ | Tension of cell membrane due to curvature | 0 | $\frac{\text{g}}{\text{s } \mu\text{m}}$ | Sensitivity analysis |

**Figure S1.**
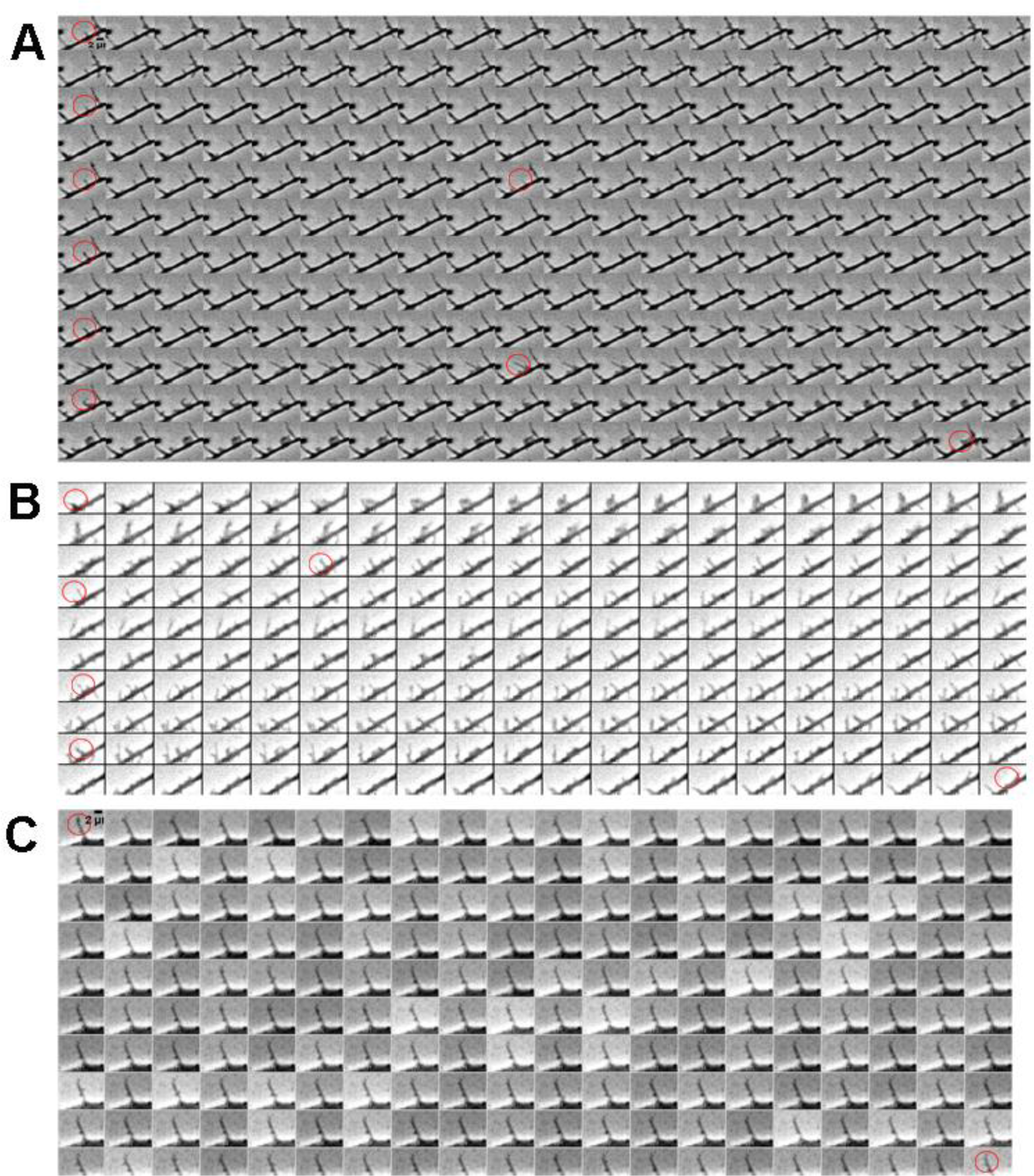
Montage of sequential phase-contrast images taken with acquisition rate 1f/s of different types of filopodial motility A. Continuously motile filopodia exhibit regular lengths fluctuations and have long lifetimes. B. Transiently motile filopodia have short lifetimes with burst dynamics. C. Non-motile filopodia have constant lengths over long periods of times.

**Figure S2.**
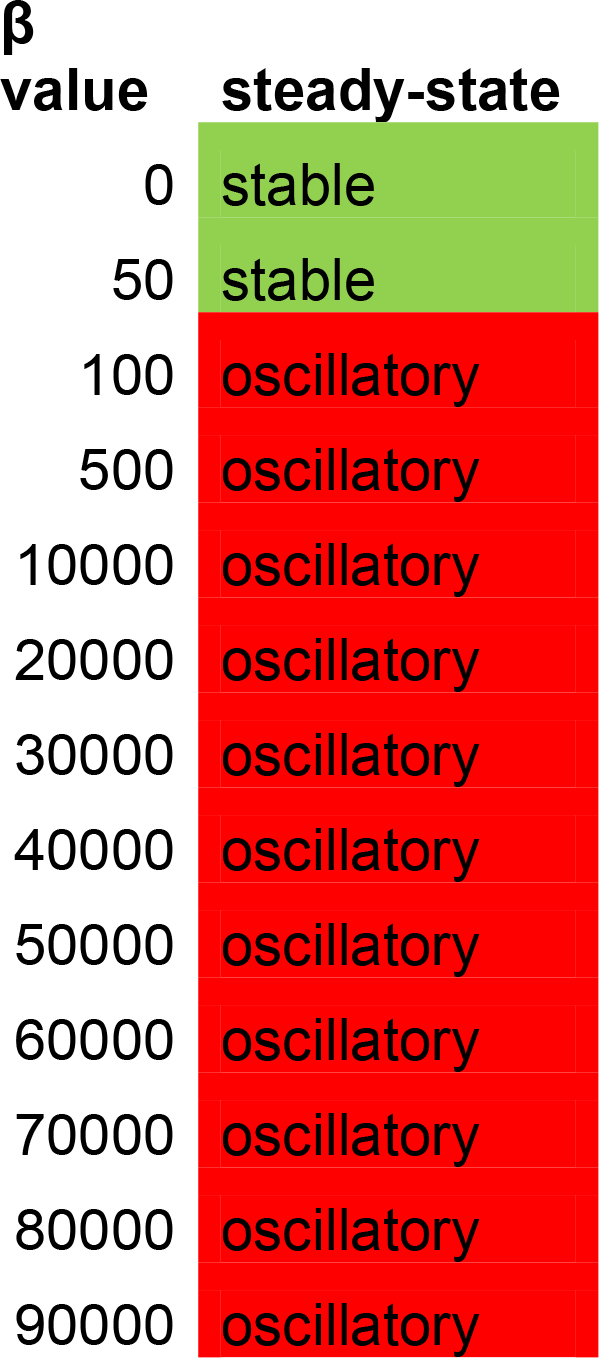
Qualitative description of resistance force parameter β significance. β values >100 result in steady-state with oscillations. Parameters used for the simulations are: k_off_=0.29, k_on_=0.27,v_p_=0.8, η =100, ζ=100, σ0 =1.3, m0=200

**Figure S3.**
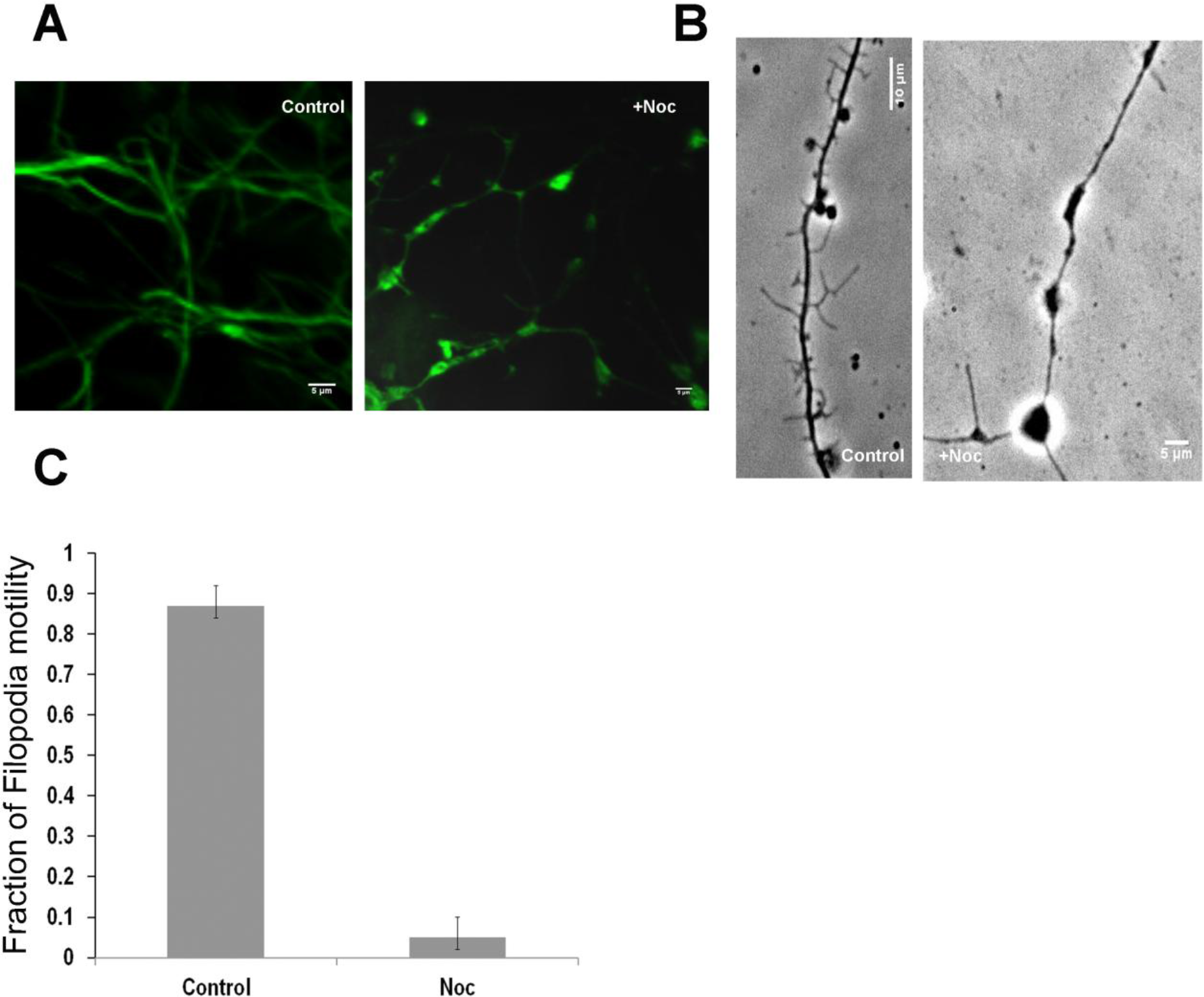
Nocodazole treatment reduces filopodia motility and number. Control neuron cultures and neurons treated with 1.6μM nocodazole after 1hr incubation at 37°C on DIV4 shown with (A) 488-Alexa tubulin staining or (B) phase-contrast microscopy. C. fraction of motile filopodia in nocodazole treated cells, t-test, P<0.05 (32 filopodia, 8 neurons).

**Figure S4.**
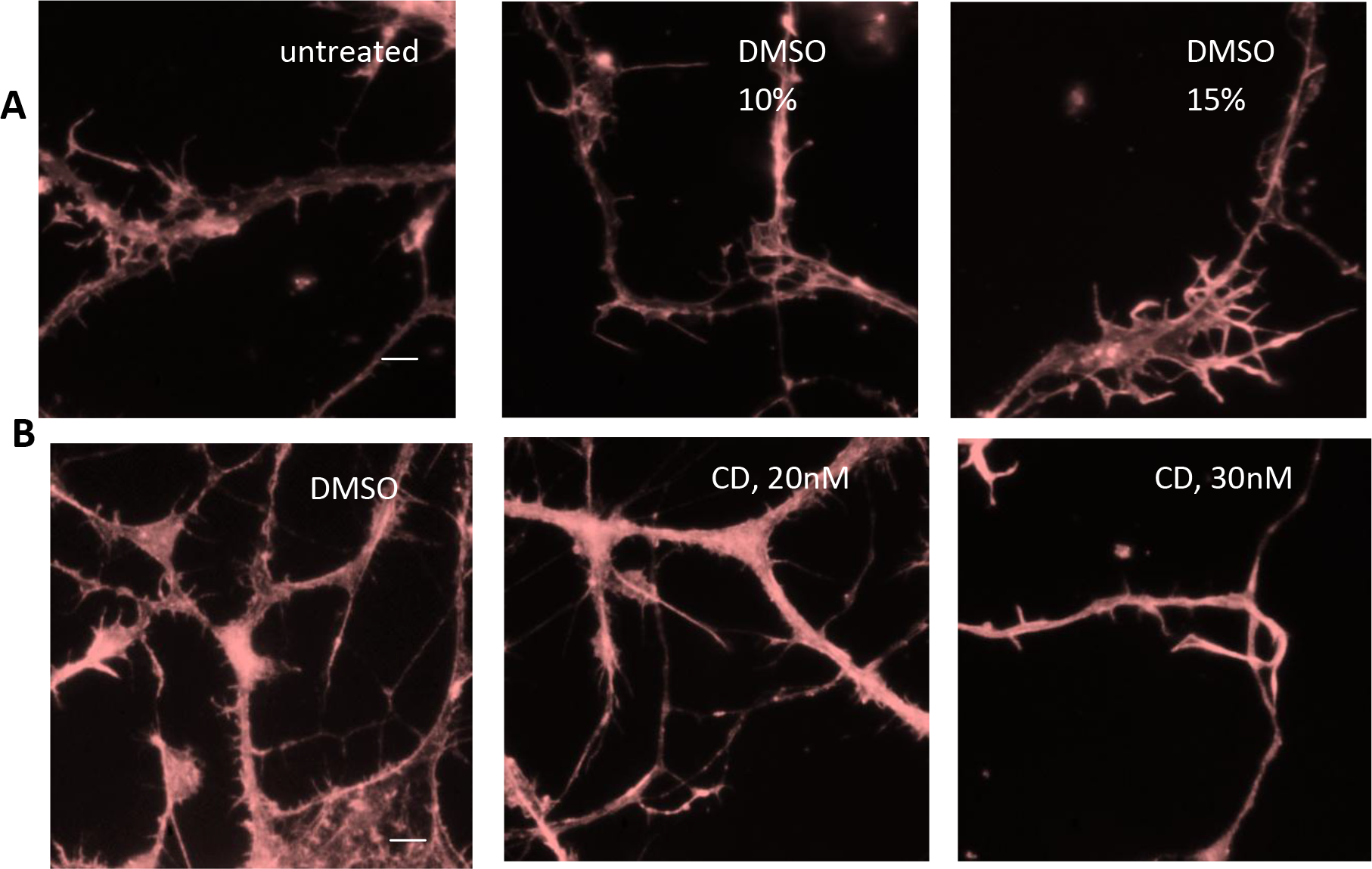
Cytochalasin D and DMSO treatments have not disrupted filamentous actin in aconcentrations used in the drug assays as shown by rhodamine staining of F-actin following drug administration. A. Control DMSO treatment did not have significant effect on density and structure of filamentous actin. B. Cytochalasin D treatment at treatment did not have significant effect on density and structure of filamentous actin at 20nM. Treatment with 30nM CD resulted in significant reduction of filopodia motility and changes in actin network compared to control. Scale bar 10μm

**Figure S5.**
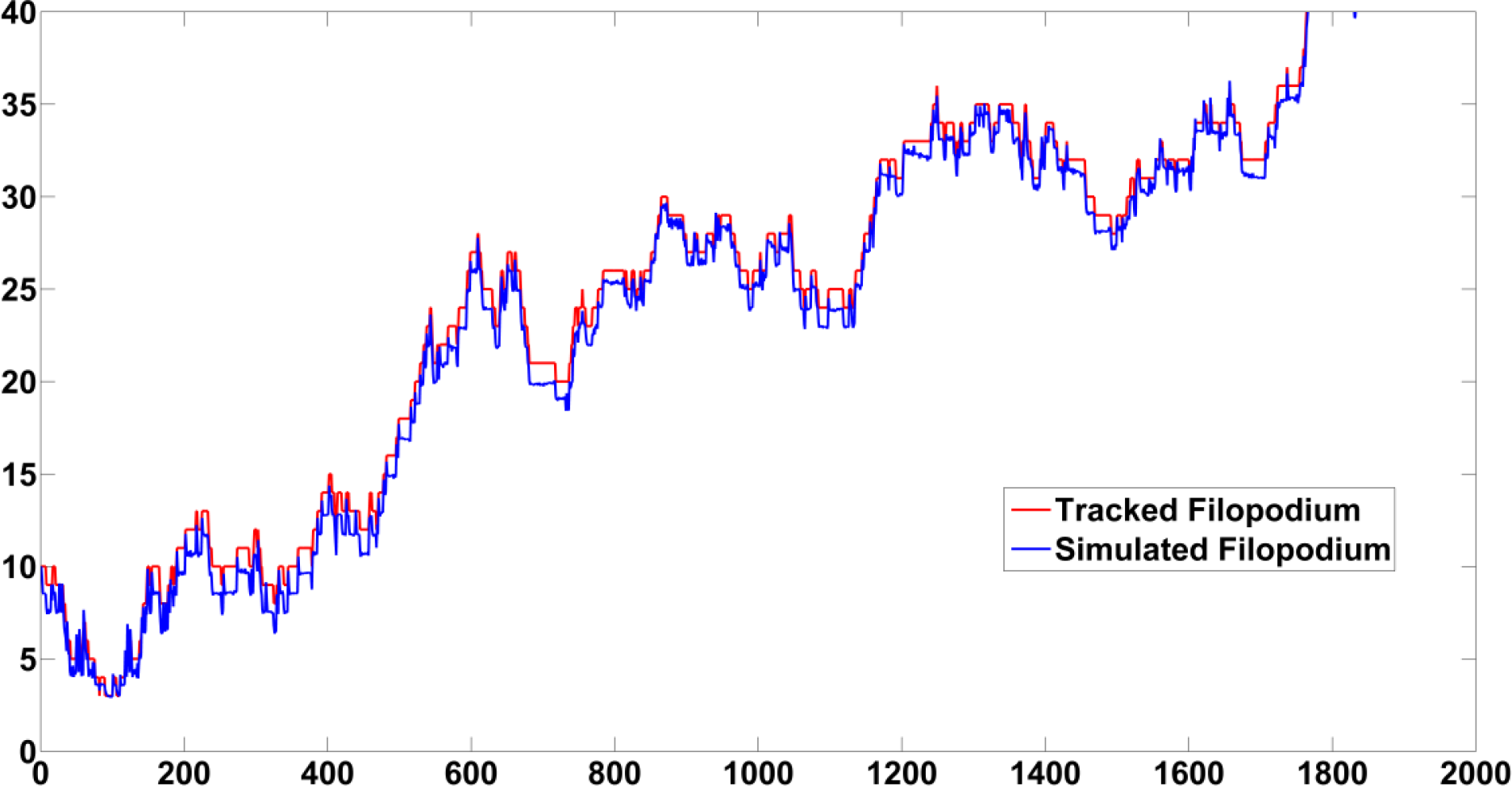
Filo tracker algorithm (red line) length measurements closely follow computer-generated binary image of filopodium dynamics simulation (blue line) with standard error E=0.88%.

**Figure S6.**
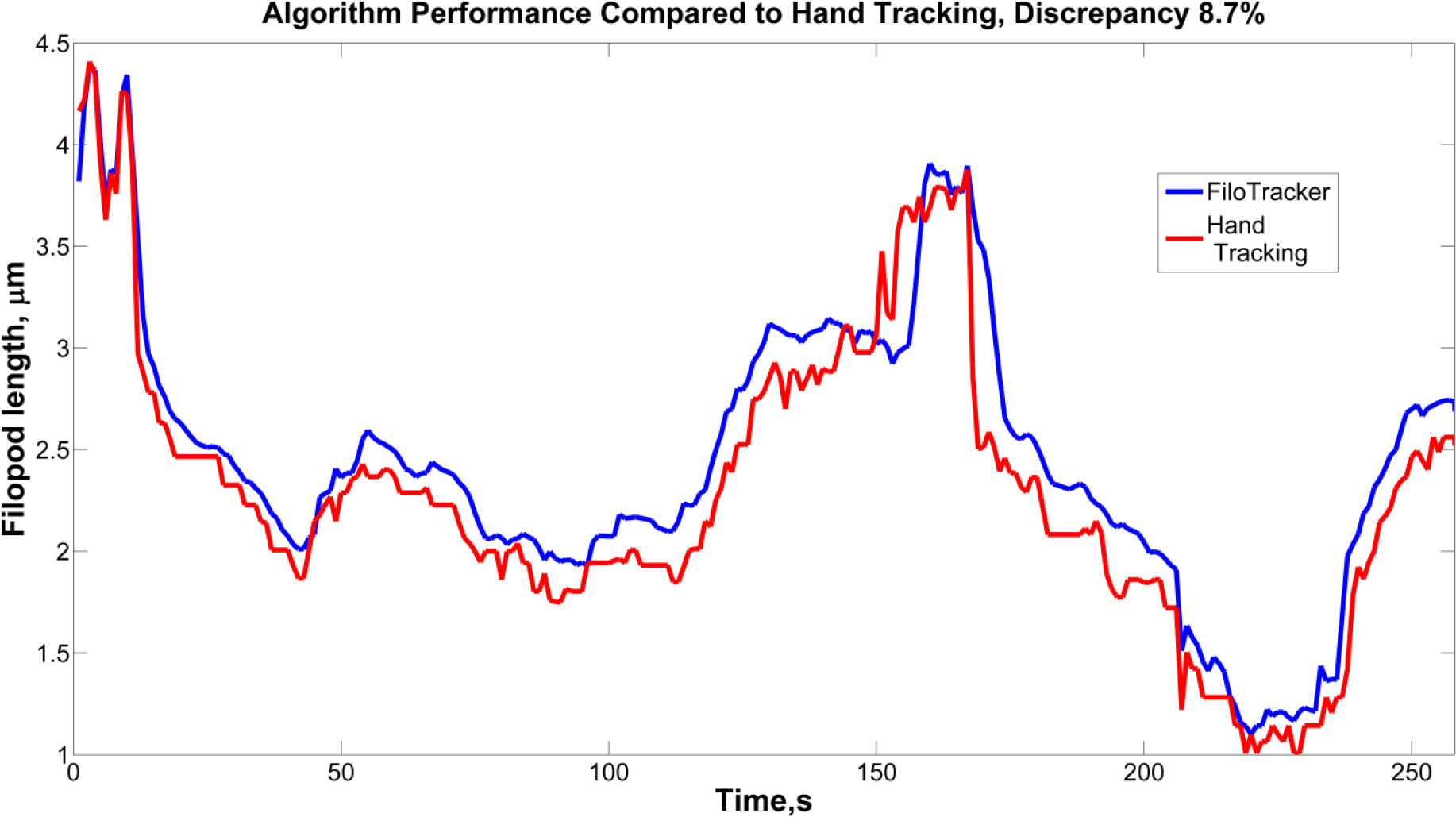
FiloTracker algorithm (blue line) length measurements output matches hand tracked filopodium lengths from live phase-contrast microscopy recording (blue line) with standard error E = 8.7%.

### 3. Legends to Supplemental Movies

*MovieS1*. Filopodium length tracked with FiloTracker Software (red dotted line), acquisition rate =4f/s, play rate 100 fr/s, rat hippocampal neuron culture, DIV4, poly-L-lysine 0.5mg/ml, bar 10μm

*MovieS2A*. - Filopodia motility after 20nM cytochalasin D treatment. rat hippocampal neuron culture, DIV4-7, poly-L-lysine 0.5mg/ml, acquisition rate =1f/s, play rate 100 fr/s,bar 10μm,

*MovieS2B*. Filopodia motility after 10nM cytochalasin D treatment, rat hippocampal neuron culture, DIV4-7, poly-L-lysine 0.5mg/ml, acquisition rate =1f/s, bar 10μm, play rate 100 fr/s

*MovieS3A*. Filopodia motility on poly-L-lysine substrate with 0.05mg/ml, rat hippocampal neuron culture, DIV4-7, acquisition rate =1f/s, bar 10μm, play rate 100 fr/s

*MovieS3B*. Reduced filopodia motility on poly-L-lysine substrate with 0.01mg/ml, rat hippocampal neuron culture, DIV4-7, acquisition rate =1f/s, play rate 100 fr/s, bar 5μm

*MovieS3C*. Reduced filopodia motility on poly-L-lysine substrate with 1.00mg/ml, rat hippocampal neuron culture, DIV4-7, acquisition rate =1f/s, play rate 100 fr/s, bar 10μm

*MovieS4*. Increased filopodia motility after treatment with 2.5μM blebbistatin, rat hippocampal neuron culture on poly-L-lysine substrate with 0.05mg/ml, DIV4-7, acquisition rate =1f/s, play rate 100 fr/s, bar 10μmSegal M.

